# Combining Anion Exchange and Size Exclusion Chromatography for Extracellular Vesicle Enrichment from Small Volumes of Human and Mouse Plasma for Quantitative Proteomics

**DOI:** 10.64898/2026.03.11.711200

**Authors:** Felicity Dunlop, Shaun Mason, Nadia Hafizi Rastabi, Sarah E. Alexander, Saeedeh Robatjazi, Jen Davis, Claire Laird, Taeyoung Kang, Suresh Mathivanan, Aaron P. Russell

**Author notes:** Corresponding Author: **Dr Felicity Dunlop,** Institute for Physical Activity and Nutrition, School of Exercise and Nutrition Sciences, Deakin University Geelong, Victoria, Australia. Email addresses. **Data availability:** A PRIDE (ProteomeXchange) dataset will be completed and publicly released upon acceptance, to ensure one-to-one alignment between the repository record and the final peer-reviewed manuscript. The record will include raw mass spectrometry files, Proteome Discoverer result files, search parameters, and sample-file mappings; the PRIDE accession and DOI will be added to the final article. For peer review, we have provided Proteome Discoverer exports in SupplementalData.xlsx and a separate SourceData.xlsx workbook containing the analyzed figure-level data (with STRING permalinks) as supplementary files.

## Abstract

Extracellular vesicles (EVs) are promising biomarkers, yet their proteomic analysis from plasma is hampered by low abundance and co-purification of contaminants (e.g., lipoproteins, platelets) and technical variability, particularly in small-volume animal models. We developed and validated a modular protocol integrating Size Exclusion Chromatography (SEC) with Strong Anion Exchange (SEC-SAX) specifically tailored for quantitative LC-MS proteomics from small starting volumes (150 μl of plasma). SEC alone successfully removed 99% of Albumin, and the SAX step significantly enriched EVs over contaminating lipoproteins. Downstream single pot solid phase enhanced (SP3) sample prep and STAGE tip solid phase extraction ensured maximum proteome depth. Critical confounding factors were objectively assessed: Platelet Factor 4 (PF4) was confirmed as a highly sensitive platelet marker, confirming the necessity of meticulous plasma preparation. Sample hemolysis impacted the plasma EV proteome data. As such, an objective measure (nanodrop spectrophotometer) of hemolysis and exclusion of hemolysed samples (heme >0.3 mg/ml) is recommended. The protocol is applicable to both human and mouse plasma as demonstrated by EV enrichment and quantification of biomarker proteins associated with neurodegenerative diseases from eight individual mouse plasma samples.

**Manuscript Highlights:**

- Developmental workflow for a quantitative SEC-SAX protocol for EV proteomics from small plasma volumes (150 μl).
- A range of variables tested including SAX beads amount, digestion buffer, digestion time, STAGE tip solid phase extraction, SAX elution buffer and sample filtration.
- The SAX step significantly enhances EV proteome depth by increasing EV purity in relation to ApoB lipoproteins.
- Shows the impact of the major confounding factors of sample hemolysis and platelet contamination on the EV proteome.
- Platelet contamination increases the number and abundance of proteins detected including known disease biomarkers and sample hemolysis is associated with proteins derived from platelet and red blood cell derived EVs.
- Platelet Factor 4 (PF4) is identified and confirmed as a sensitive marker for platelet contamination.
- Applicable to both human and mouse plasma.

## 1 Introduction

Extracellular Vesicles (EVs) are nanosized, lipid-bilayer vesicles released by nearly all cells, encapsulating protein and RNA cargo that reflects their cell of origin (Du et al., 2023; Jeppesen et al., 2024; Manno et al., 2024; van Niel et al., 2018). Functioning as a critical intercellular messaging system, EVs are attractive biomarkers, especially for conditions like neurological diseases and malignancies, as they can be accessed via a relatively non-invasive liquid biopsy (blood) (Malaguarnera & Cabrera-Pastor, 2024; Rayamajhi et al., 2024; Soleymani et al., 2023). Applying liquid chromatography-mass spectrometry (LC-MS)-based proteomics to the plasma EV proteome holds potential to generate quantitative data on thousands of proteins, significantly enhancing our understanding of the role EVs play in health and disease (Garza et al., 2024; Greening et al., 2024; Rai et al., 2025). However, the effective analysis of plasma EVs is challenged by low EV abundance relative to other plasma components. Major contaminants such as lipoproteins, albumin, and immunoglobulins co-purify with EVs (Sodar et al., 2016; Dmitry Ter-Ovanesyan et al., 2023a; Veerman et al., 2021). Furthermore, hemolysis and platelet contamination are recognized as significant confounding factors in blood-derived EV studies (Coumans et al., 2017; Lucien et al., 2023). These contaminants negatively impact the purity of the EV preparation, compromising the accuracy of any identified protein signature.

Guidelines for blood sample collection and plasma preparation include numerous recommendations, such as needle gauge, tube transport, centrifugation temperature and timing and speed to remove red blood cells and platelets, while avoiding hemolysis, platelet activation and platelet EV release (Coumans et al., 2017; Thery et al., 2006). Despite these recommendations, the full impact of these factors is often under-quantified in proteomics studies (Holcar et al., 2021; Korff et al., 2025; Rios de Los Rios Resendiz et al., 2025). The Methodological Guidelines to Study Extracellular Vesicles recommends that EVs from hemolysed samples should not be used (Coumans et al., 2017; Lucien et al., 2023; Thery et al., 2006). However, the MIBlood-EV lists visual inspection as a qualitative method for the assessment of plasma hemolysis despite being inaccurate and highly subjective with hemolysis not visually apparent until 0.3 g/l heme (Gislefoss et al., 2021)(Lucien et al., 2023; Wan Azman et al., 2019). Additionally, the impact of sample hemolysis on EV enrichment and the EV proteomics signature is largely unknown (Rios de Los Rios Resendiz et al., 2025).

Standardizing upstream sample preparation could potentially improve the reproducibility of plasma-derived EV data. However, the extent to which plasma sample preparation can be standardised is potentially limited by differences in individual laboratory protocols and the generic nature of plasma biobank collections (Lucien et al., 2023; Venturella et al., 2019). Additionally, guidelines are largely developed with plasma samples from humans in mind, which may not be feasible when dealing with small plasma volumes from small animals. Furthermore, the lack of a quality control metrics, specifically for platelet contamination of LC-MS based plasma EV proteomics data, raises concerns as to the origin of the proteomics signature identified.

The challenge of EV purification from plasma persists. Size Exclusion Chromatography (SEC) is a common technique used to enrich plasma EVs by removing a large proportion of soluble plasma proteins (Dong et al., 2020; Sidhom et al., 2020; Stranska et al., 2018). Despite its effectiveness against soluble proteins, SEC also enriches lipoproteins (such as VLDL) due to the significant overlap in size between small EVs and lipoprotein subclasses (Sódar et al., 2016; Veerman et al., 2021). Combining approaches that leverage differences in physical and biological properties between EVs and the free plasma proteins and lipoproteins is effective for enrichment of the plasma EV population for proteomics (Stam et al., 2021; Su et al., 2024; D. Ter-Ovanesyan et al., 2023b; Van Deun et al., 2020). Surprisingly, EV enrichment techniques continue to be published that claim to sufficiently enrich EV from blood plasma/serum in a single step (Ji et al., 2021; Khanabdali et al., 2024; Wu et al., 2025). However, the reliance of these methods on highly sensitive techniques like Data Independent Acquisition (DIA) or Field Asymmetric Ion Mobility Spectrometry (FAIMS) suggests that the extent of EV enrichment may be questionable, as it indicates a need for increased MS sensitivity to overcome residual contaminants. (Ji et al., 2021; Khanabdali et al., 2024; Lattmann et al., 2024; Vanderboom et al., 2021; Wu et al., 2025).

There is a clear need for proteomic evaluation of EV purity, defined as the ratio of EV to non-EV components, and for the systematic identification of confounding factors, such as lipoproteins, hemolysis and platelets. Therefore, the aim of this study was to develop and optimise a robust, quantitative protocol for enriching small EVs from human and mouse plasma that is suitable for LC-MS-based proteomics. Throughout protocol development, emphasis was placed on objective quality-control measures to detect and account for key confounders. This approach ensured the generation of high-purity EV proteomes suitable for reliable quantification and accurate biomarker discovery.

## Materials and Methods

For clarity, the Results section begins with a description and summary table of the 11 experiments that informed the development and testing of the SEC-SAX protocol; this overview accompanies the full detailed results presented throughout the section.

### 2.2 Blood collection and plasma preparation

#### 2.2.1 Human plasma

Blood was taken from a healthy male donor in the fasted state. A 21-gauge needle (cat#AN*2125R1, Terumo, Tokyo, Japan) was used to collect blood from the antecubital vein using a vacutainer tube containing 7.2 mg K2-EDTA (cat #367839, Becton Dickinson, Franklin Lakes, NJ). Blood was collected directly from the needle into the vacutainer tube using a Greiner Holdex tube holder (cat#450263_CT, Greiner Bio-One, Kremsmünster, Austria) without the use of a syringe or aspirating device, which may cause haemolysis. Plasma was prepared without filtration but otherwise essentially as described by (Dong et al., 2020). Briefly, whole blood was separated via centrifugation for 10 min at 1000 g, 22°C, acceleration and deceleration setting 1 (acc/dec 1) (Multifuge XR4Pro centrifuge with XT-1000 rotor). The plasma was gently transferred to fresh 15 ml tubes using a P1000 pipette and centrifuged again for 15 min at 2500 g, 22°C, acc/dec 1. The plasma was transferred to fresh 15 ml tubes and the 2500 g spin repeated. Finally, the plasma was gently transferred to fresh 50 ml tube to generate a pool of plasma and then aliquoted as 550 µl or 1100 µl and stored at-80°C. Plasma was thawed on ice and briefly hand warmed with mixing by inversion to dissolve cryo-precipitates. Plasma was precleared to remove larger vesicles by centrifugation at 10000 g, 20 minutes, 4°C (Thermo Fisher Heraeus Megafuge 8R with 75005715 rotor) and the supernatant transferred to a new 1.5 ml tube. Precleared plasma was kept on ice until processed by Size Exclusion Chromatography (SEC). Deakin University Human Research Ethics Committee (DUHREC) approved project 2018-388.

#### 2.2.2 Mouse plasma

Mouse plasma was collected by cardiac puncture using a 1 ml syringe (Terumo, cat#SS+01T) primed with 0.5M EDTA (Invitrogen, cat#J15694.AP) fitted with a 21-gauge needle (Terumo, cat#AN*2138R1). Blood was transferred to Multivette 600 µl KE3 tubes (Sarstedt, cat#15.1671.100) and prepared using the same centrifugation protocol (1000 g 10 min, 2500 g 15 min x2) as for human plasma but at 1.5 ml tube scale. The Heraeus Megafuge 8R centrifuge with 75005715 rotor (Thermo Fisher) was used with the soft acceleration and deceleration setting enabled to minimise shear stress and plasma activation. The heme content was determined by spectroscopy (Section 2.9) prior to storage at-80°C and the mass of plasma determined and recorded for each individual mouse aliquot by the difference in mass between the 1.5 ml tube with and without plasma. Deakin University Animal Ethics Committee approved project 2020/AE025980.

### 2.3 Size exclusion chromatography (SEC)

#### 2.3.1 Automated size exclusion chromatography using an automatic fraction collector (AFC)

The AFC-V1 was used to process plasma by SEC for the first two experiments (Sections 3.1-3.2). Briefly, phosphate buffered saline (Sigma-aldrich, cat#P4417-100 TAB) was made up fresh with MilliQ water using dedicated SEC glassware and filtered with a 0.2 µm syringe filter (Whatman Uniflo, cat# 9914-2502) prior to use. qEVoriginal 35 nm Legacy resin columns (Izon Science, cat#SP5) were equilibrated at room temperature and the automatic fraction collector (AFC-V1) used to process 500 µl of plasma with default void volume of 2.8 ml discarded before collection of 0.5 ml fractions. Columns were run with a total of 10 ml of PBS before cleaning with 10 ml 0.5 M filtered NaOH and flushing with 30 ml PBS. Columns were reused a total of 5 times before discarding according to the manufacturer’s instructions. Fractions were kept on wet ice until protein content was measured by absorbance at 280nm (Section 2.9). Samples were stored as individual fractions (Experiment one, Section 3.1) or pooled and concentrated fractions (Experiment two, Section 3.2) stored at - 80°C until western blot analysis (Section 2.10).

#### 2.3.2 Manual size exclusion chromatography

Plasma was processed manually for experiment three onwards (Sections 3.3-3.11).

##### 2.3.2.1 qEV original 35 nm Legacy resin (Izon Science, cat#SP5)

Manual processing of plasma by SEC (Izon Science, cat#SP5) was simply performed by accurately pipetting the volume of buffer required using analytical pipettes (Gilson) and waiting for the buffer to completely drip through. A P10 ml pipette was used for flush and buffer volumes of 10 ml-30 ml. A P1000 µl pipette was used for pipetting the void of 2.7 or 2.8 ml as two lots of 1 ml and then the remaining 0.7 or 0.8 ml. A P1000 was also used for pipetting individual 0.5 ml fractions with the buffer allowed to drip into a collection tube until no more liquid remained and then the next fraction collected in the next tube by the addition of another 0.5 ml of buffer. For experiments 4 and 5 (Sections 3.4 and 3.5), the 2.7 ml void was used and pooled fractions 1-4 eluted directly into the centrifugal filter by addition of 2 ml of PBS to the SEC column using a P1000 twice instead of individual fractions.

Preparation of EV from 500 µl of human plasma was performed the same as for the large scale 4 ml but as a single lot of 500 µl. Fractions 1-4 (total 2 ml) were pooled and concentrated down to a final volume of 170 µl and stored as 2 aliquots of 85 µl at-80 °C until analysis by Western blotting.

##### 2.3.2.2 qEVsingle 35 nm Gen 2.0 resin (Izon Science, cat#ICS35)

For the first use of the smaller qEV single columns (Izon Science, cat#ICS35) which take 150 µl of plasma the void volume of 700 µl was used and increasing amounts of SEC fractions were pooled for SAX enrichment and proteomics analysis (Section 3.5). The storage buffer was allowed to run out of the column and the column was equilibrated with 3 column volumes of Bis-Tris Propane SEC buffer (3 ml each = 9 ml) (50 mM BTP (Sigma-Aldrich, cat#B9410), 75mM NaCl (Sigma-Aldrich, cat#S3014), pH 6.3). Once the column had stopped dripping and the frit was just dry 150 µl of precleared plasma was pipetted directly on the top of the frit using a P200. Once the plasma had completely run into the column (as indicated by a dry frit and the column no longer dripping) 550 µl of BTP SEC buffer was added to the top of the column for a total void volume of 700 µl (150 µl plasma + 550 µl buffer). The void volume was allowed to run through and was discarded. Either 4, 5 or 6 x 170 µl fractions were collected as 680 µl, 850µl or 1020 µl directly into 1.5 ml Eppendorf tubes containing 20 µl SAX beads already equilibrated twice with 600 µl each BTP Bead buffer (50 mM Bis Tris Propane, 150 mM NaCl, pH 6.5).

For the remaining experiments (Sections 3.6-3.11), the void of 700 µl (150 µl plasma + 550 µl BTP) and the SEC eluted in a single 680 µl fraction. qEV single columns were discarded after a single use.

### 2.4 Centrifugal filtration

Amicon Ultra Centrifugal Filters, 100 kDa MWCO (cat#UFC910008) were used to concentrate material eluted from the SEC for experiments one to five (Sections 3.1-3.5). Centrifugal filters were centrifuged at 4000 g at 4⁰C (Multifuge XR4Pro centrifuge with XT-1000 rotor) until the indicated volume for each experiment and the concentrate removed using a P200 pipette.

### 2.5 Ultracentrifugation

The Ultracentrifugation protocol was adapted from Current Protocols in Cell Biology Basic protocol 2 (Thery et al., 2006) for experiment two (Section 3.2). A 1.5 ml Eppendorf tube was used to dilute 500 µl of plasma with 500 µl of 0.2 µm filtered PBS and centrifuged 2 000 g, 30 minutes at 4⁰C (Heraeus Megafuge 8R centrifuge with 75005715 rotor, Thermo Fisher). The supernatant was transferred to a fresh 1.5 ml tube using a P200 and centrifuged again at 12 000 g, 45 minutes at 4⁰C. The supernatant was transferred to an ultracentrifuge tube (Beckman Coulter, cat # 362305) and centrifuged 110 000 g, 2 hours at 4⁰C using an Ultracentrifuge (Optima Max-XP and a fixed angle TLA-110 rotor, Beckman Coulter). The supernatant was discarded and the EV pellet resuspended in 4 ml PBS and centrifuged at 100 000 g for 70 minutes at 4⁰C. Removal of the supernatant, resuspension of the pellet in 4 ml PBS and centrifugation at 100 000 g for 70 minutes at 4⁰C was repeated. The final EV pellet was resuspended in 100 µl of PBS and stored at-80⁰C until western blotting (Section 2.10).

### 2.6 EV Concentration kit (Izon Science)

For Experiment Two (Section 3.2) pooled SEC fractions 1-4 from 35 nm qEVoriginal (legacy) was eluted directly into a 2 ml tube containing 50 µl of EV concentration kit (Izon Science, cat# RCT02). Beads were incubated overnight on a suspension mixer (Ratek Instruments, cat# RSM6) at 4⁰C and then the beads collected by centrifugation at 16 000 g 10 minutes at 4⁰C. The supernatant was discarded and the beads heated at 95⁰C for 7 minutes in 55 µl NuPage LDS sample loading buffer (Thermo Fisher, cat#NP0007) + β-mecaptoethanol (Bio-Rad, cat#1610710). The beads were pelleted by centrifugation at 16000 g 10 minutes, the supernatant transferred to a fresh tube and stored at-80⁰C for subsequent analysis by western blotting (Section 2.10). For Experiment Five (Section 3.5) 150 µl of concentrated SEC material (SEC+CF) diluted with an equal volume of PBS and incubated overnight with 50 µl of EV concentration kit as for experiment 2. Beads were eluted in 100 µl 2% SDS 100 mM HEPES for LC-MS analysis (Section 2.13).

### 2.7 Immunoaffinity capture (CD9, CD63, CD81 magnetic beads)

For Experiment Five (Section 3.5) Immunoaffinity capture (IAC) of EVs from biological fluid using magnetic beads was adapted from the plasma EV RNA purification protocol (Heinzelman, 2018). Proportional volumes of CD9 (20µl) (Thermo Fisher, cat #10614D), CD63 (30 µl) (Thermo Fisher, cat #10606D), and CD81 (20 µl) (Thermo Fisher, cat #10616D), beads based on package size were combined and washed once in 600 µl PBS. Concentrated SEC material (150 µl) was diluted up to 300 µl in PBS and incubated overnight at 4°C on a suspension mixer (Ratek Instruments, cat# RSM6). Beads were washed 3 times with PBS prior to elution in 100 µl 2% SDS 100 mM HEPES for LC-MS analysis (Section 2.13).

### 2.8 Anion Exchange (SAX magnetic beads)

For Experiment Five onwards (Sections 3.5-3.11), Strong Anion Exchange (SAX) magnetic bead purification was adapted from EV purification from plasma for proteomics (Wu et al., 2025). Individual aliquots (20 µl, except where indicated in Sections 3.5 and 3.6) of SAX beads (ReSyn Biosciences, MR-SAX010) were equilibrated twice with 600 µl each BTP Bead buffer (50 mM Bis Tris Propane (Merck, B9410) 150 mM NaCl (Sigma-Aldrich, S3014), pH 6.5) for 5 minutes each, 4°C on a suspension mixer (Ratek Instruments, cat# RSM6). A magnetic rack (Thermo Fisher cat#12321D) was used to capture the beads for washing and removal of the bead buffer. The bead buffer was removed from the equilibrated beads immediately prior to SEC elution to prevent beads drying and the SEC columns eluted directly into the tubes containing beads.

For the filtration experiment (Section 3.9) the 680 µl of SEC eluant was first eluted in a separate 1.5 ml tube and then filtered through a 0.2 µm PES syringe filter (Nalgene, cat#Z741696) using a 1 ml syringe. To minimise EV loss the filter was flushed with 680 µl of BTP Bead buffer and combined with the filtered SEC eluant for binding to the SAX beads.

Phosphatase and protease inhibitor (Thermo Fisher, cat#78446) was added to the SEC eluant and incubated overnight at 4°C on a suspension mixer (Ratek Instruments, cat# RSM6). SAX beads were washed three times (900 µl) in ice cold Bis-Tris Propane bead buffer and then the beads resuspended and eluted in 200 µl 2% SDS, 100 mM HEPES pH 7.5 (15 minutes 95°C) for LC-MS analysis (Section 2.13).

### 2.9 Spectroscopy

#### 2.9.1 Heme content

The level of hemolysis of precleared plasma was measured using the Nanodrop One spectrometer (ThermoFisher) using the custom Oxy-hemoglobin method with 2µl of plasma and water as the blank. A cutoff of >0.3 was used to exclude hemolysed samples (Shah et al., 2016).

#### 2.9.2 Protein content

The protein content of the SEC fractions was measured using the Nanodrop spectrometer by measuring absorbance at 280 nm. For each measurement, 2 µl of the fraction was used, with PBS buffer used in the SEC as the blank.

### 2.10 Western blotting

Samples were resolved on 4-15% Criterion Stain free gels (Biorad, cat# 5678085) at 200 Volts, 40 minutes with Precision Plus Protein All Blue standard 4 µl (Biorad, cat#161-0393). Proteins were transferred to a PVDF membrane by Western blotting at 100 volts for 1 hour. Membranes were blocked in 5% skim milk in PBS at room temperature for 1 hour before overnight incubation with primary antibodies Syntenin (Abcam, cat# ab133267), CD63 (Abcam, cat#ab134045), CD63 (Abcam, cat#ab8219), Albumin (Cell Signalling Technology, cat#4929S), Calnexin (Cell Signalling Technology, cat#2679S) ApoA1 (Santa Cruz, cat# SC-376818). Blots were washed 3 times in PBS + 0.05% Tween 20 for 10 minutes each. The corresponding fluorescent secondary antibodies (Cell Signalling Technology, anti-mouse 680 nm (cat# 5470S), anti-mouse 800nm (cat#5257S), anti-rabbit 800 mm (cat#5151S) were incubated at room temperature for 1 hour and the blots were washed twice in PBST and once in PBS. The blots were then simultaneously imaged at 680 nm and 800 nm using the infrared imaging system (Li-COR Biosciences, Lincoln, NE, USA).

### 2.11 Electron Microscopy

For Experiment Three (Section 3.3), the protein content of the concentrated SEC material was first determined using the Pierce BCA protein assay (Thermo Fisher, cat# 23225) and adjusted to 0.2 mg/ml with sterile filtered PBS. 400-mesh formvar-carbon coated copper grids (ProSciTech, Australia) were glow discharged for 60 s using Carbon Coater K950X with K350 attachment (Quorum Emitech, UK). Samples were applied onto copper grids and allowed to absorb for 30 s. Excess liquid was carefully removed using filter paper and immediately after, EVs were stained with 2% (w/v) uranyl acetate (ProSciTech, cat#C079) for 10 s and dried again using filter paper. JEM-2100 electron microscope (JEOL, Japan) equipped with AMT NanoSprint15 Mk-II camera (AMT TEM Imaging Systems, USA) was used for visualisation at 200 kV with 30,000× magnification.

### 2.12 Tunable resistive pulse sensing: particle size and concentration

For Experiment Three (Section 3.3), the size and concentration of the particles in the pooled SEC eluted EV fractions 1-4 (prior to concentration by centrifugal filter) was determined using a qNano (Izon Science) using the NP 80 (80 nm) nanopores with the qNano reagent kit (Izon Science, cat#RK3). Nanopores were coated with the Izon coating solution in measurement electrolyte (ME) for 5 minutes at ≥18 mbar and 2X measurement electrolyte (2ME) was used for subsequent measurements to increase the signal of the small particles. A concentration standard curve method was used to determine the concentration of the particles in the pooled SEC fractions instead of a pressure dependent method as the NP80 nanopores became blocked with increasing pressures. The particle rate (particles passing through the pore per minute) was proportional to the concentration of the 60 nm calibration beads so a minimum of a 3-point standard curve of concentrations over the range of 5 x10^9^ - 2 x10^10^ of particles per ml was generated for each nanopore. The SEC fractions were diluted to produce a particle rate within this linear range and the particle rate of the SEC material converted to a concentration in particles per ml from the standard curve. Due to the highly variable and unstable nature of the NP80 nanopores the standard curve measurements were considered valid if duplicate samples of the top calibration point produced the same particle rate and 2-fold dilution resulted in the particle rate halving. Sample measurements were only considered valid if the RMS<10, the current was stable with a drift of <10 pA and particle rate was linear. Where possible the samples were measured in duplicate while the nanopore remained stable.

### 2.13 Proteomics analysis

#### 2.13.1 Sample Preparation for Proteomics

Samples were prepared for analysis by LC-MS based proteomics after lysis to release and solubilise the EV proteins. Samples were simultaneously reduced and alkylated and then prepared by single spot solid phase enhanced (SP3) (Hughes et al., 2019). For Experiments 3 and 4, the protein content was first determined by BCA protein assay and 10 µg SEC material (pooled fractions 1-4 concentrated by centrifugal filtration) was diluted to a final volume of 90 µl 2% SDS (Sigma-Aldrich, cat#05030), 100 mM HEPES (Sigma-Aldrich, cat#SRE0065) and prepared for LC-MS analysis as for samples prepared with the additional bead-based EV enrichment processes. For Experiment Five, EV enriched samples using the additional SAX, IAC and CK enrichment steps were eluted in 100 µl 2% SDS, 100 mM HEPES by boiling at 95°C for 15 minutes. The samples were cooled on ice and then placed on the magnetic rack for 2 minutes before the eluted proteins were transferred to a fresh 1.5ml tube and the spent beads discarded. The non-magnetic CK beads were removed by centrifugation 16,000 g, 10 minutes at 4⁰C and the supernatant transferred to a fresh tube. For Experiment 6 onwards the SAX beads were eluted in 200 µl 2% SDS, 100 mM HEPES to aid in recovery of the EV proteins as the SAX beads stick to the tubes when combined with human SEC material. Disulfide bond reduction and alkylation was done by separate addition of TCEP (Sigma-Alrich, cat #646547) and IAA (Sigma-Alrich, cat #I6125) to a final concentration of 9 mM and 40 mM respectively, and subsequent heating the samples at 95⁰C for 15 minutes in the dark. Five mg of each Sera-Mag Hydrophilic and Hydrophobic magnetic SpeedBeads (Cytiva, cat# 45152105050250, cat#65152105050250) were combined and washed with 1 ml LC-MS water (cat #1.15333) on a magnetic rack. Washed beads were resuspended in 1 ml LC-MS water and 20 µl added to each tube of eluted proteins. Binding of the proteins to the beads was induced by addition of an equal volume (120 µl or 240 µl) of absolute ethanol (Sigma-Alrich, cat #459836) and mixed using a Thermomixer (Eppendorf, Macquarie Park, Australia) at 1600 rpm for 8 minutes. The bead-bound proteins were washed three times with 80 % ethanol before resuspension in 60 µl digestion buffer (10% TFE (Sigma-Aldrich, cat#T63002), 100 mM HEPES pH 7.3) and 0.1 µg trypsin (Thermo Fisher, cat #90305) and incubated for 18 hours at 37⁰C, 1600 rpm using a Thermomixer.

STAGE tips: Overnight trypsin digests were desalted using STAGE tips prepared from 2 layers of SDP-RPS membrane (Empore, cat # 2341). The STAGE tips were first equilibrated in 50 µl each of acetonitrile (Sigma-Aldrich, cat #1.00029), 0.2 % TFA (Sigma-Aldrich, cat# T0699) in 30% methanol (Sigma-Alrich, cat#1.06035), and then 0.2% TFA by centrifugation at 500 g for 3 minutes each. Overnight trypsin digests were placed on the magnetic rack. The supernatant containing the digested peptides were pipetted on top of equilibrated STAGE tip containing 150 µl 1 % TFA and centrifuged at 1000 g for 8 minutes. Tips were washed twice with 100 µl 0.2% TFA and once with 40 µl 1% TFA in 90% isopropanol by centrifugation at 1000 g for 5 minutes. Peptides were eluted into a new 1.5 ml tube with 60 µl 5 % ammonium hydroxide (Sigma-Alrich, cat#221228) in 80 % acetonitrile by centrifugation at 1000 g for 3 minutes. Eluted peptides were dried in a CentriVap (Labconco, Kansas, USA) at 45⁰C for 45-60 minutes until dry and stored at 4⁰C (short term) or-80°C (long term). Peptides were resuspended in 10 µl (Experiments 3.5-3.11) or 20 µl (Experiments 3.3-3.5) in loading buffer (2% acetonitrile, 0.1% TFA) for liquid chromatography mass spectrometry analysis (LC-MS).

#### 2.13.2 Liquid chromatography mass spectrometry

Samples were analyzed using an UltiMate 3000 RSLCnano (ThermoFisher Scientific) linked to an Orbitrap Exploris™ 240 Mass Spectrometer (ThermoFisher Scientific). Six or nine microlitres of sample (equal peptide volume) was loaded onto a PepMap 100 C18 (20 mm x 0.1mm, 5µm) nanoViper trap column for 3 mins at a flow rate of 15 µL/min with 2% (v/v) acetonitrile, 0.1% (v/v) TFA, and then resolved on a PepMap Neo C18 (75 µm x 500 mm, 2µm) nanoViper analytical column at a flow rate of 250 nL/min. Mobile phases consisted of 0.1% (v/v) Formic acid (A) and 80% (v/v) Acetonitrile/0.1% (v/v) Formic acid (B). Gradient conditions were: 0-2 min 2.5% B, 2-2.1 min 8% B, 2.1-122 min 32% B, 122-125 min 45% B, 125-133 min 99% B, 133-150 min 2.5% B. The column compartment was set at 50°C for heated trap elute. Ion source conditions were: NSI type, static spray voltage, 2000V positive ion, 1500V negative ion, 275°C ion transfer tube temperature. MS1 data were acquired over 350-1200 m/z with an orbitrap resolution of 120000, RF lens 70%, and positive polarity. Dynamic exclusion was for 20s, with mass tolerances of 10 ppm. Data dependent acquisition used a 3 s cycle time with HCD collision energy set at 30%. MS2 spectra were acquired with a fixed first m/z of 120 with an orbitrap resolution of 15 000.

#### 2.13.3 Data Analysis and software

Data was processed using Thermo Proteome Discover (v.3.1) with Sequest HT as the database search algorithm against the *Homo sapiens* for human samples (Experiments 3.3-3.10) and *Mus musculus* (Experiment 3.11) FASTA file (Uniprot). Input data to Sequest HT were as follows: Maximum missed tryptic cleavages 2, precursor mass tolerance 10 ppm, fragment mass tolerance 0.02 Da. Dynamic modifications were oxidation (methionine), and static modifications were carbamidomethylation (cysteine). The processing workflow also had the Percolator node used, with strict target FDR set at 0.01 and relaxed FDR set at 0.05. The target/decoy strategy was concatenated with validation based on the q-value. Data Normalization was not used (with the exception of Section 3.3) as the starting volume of plasma was used as the comparison point. Data normalization was based on total peptide amount for Section 3.3 as an equal protein amount was used for each sample (10 µg). Protein abundances were calculated on the basis of summed peptide abundances of unique and razor peptides and no imputation was used. Protein ratios were calculated using protein abundances and Hypothesis testing was based on ANOVA (individual protein) calculations. Fold-change individual protein ratios of >1.5 (Log2=0.58) and ANOVA p-value <0.05 were used as statistical cutoffs to determine significance. Heatmaps and PCA plots were exported directly from Proteome Discoverer. Additional larger labels were added to the PCA plot using Microsoft Powerpoint. Volcano plots were generated using VolcaNoseR (https://huygens.science.uva.nl/VolcaNoseR/)(Goedhart & Luijsterburg, 2020). Numerical data was exported from Proteome Discoverer as Microsoft Excel spreadsheets.

Quantified proteins were defined as those with an abundance >0. Grouped abundances, calculated by Proteome Discoverer (average protein abundance for a protein in a defined sample group) was used to identify the number of quantified proteins for a group. STRING analysis (https://string-db.org/) was used to identify the number of proteins categorised as Cellular Component Extracellular Vesicles GO:1903561 using the unique accession numbers for human samples. Comparison of the unique accession numbers with the “real-vesicular proteins” as a stringent measure of the number of EV markers quantified (Choi et al., 2020). Intrasample EV purity ratios were calculated for CD9, CD81 and Syntenin-1 abundance in relation to the contaminant proteins Albumin and ApoB in Microsoft Excel. Additional statistical analysis and graphs were generated using GraphPad Prism. Method images were generated with BioRender. SRplot was used for visualisation of STRING GO: analysis (Tang et al., 2023).

A detailed step-by-step version of the final SEC-SAX protocol is included as Appendix A.

## 3.0 Results

The development and testing of the SEC-SAX protocol required 11 experiments. The first 3 experiments (**3.1-3.3**) established the protocol for Size Exclusion Chromatography including analysis for: total protein content measured via spectrometry at an absorbance at 280 nm, western blot for EV and contaminating proteins, particle concentration and size by TRPS and quantitative proteomics analysis.

Experiments **3.4** and **3.5** detail the screening of EV enrichment techniques that, in combination with SEC, increase EV purity when determined via proteomics analysis and the subsequent removal of redundant steps respectively.

Experiments **3.6-3.8** optimise the SAX bead amount, SAX elution buffer, and the necessity of the solid phase extraction post digestion desalting step respectively.

Experiments **3.9-3.10** test the impact of confounding factors including, platelet contamination and sample hemolysis in human plasma. They also investigate the suitability of a filtration step for inclusion in the SEC-SAX protocol.

Experiment **3.11** tests the SEC-SAX protocol in a cohort of 8 individual mouse plasma samples. Refer to Table 1 for a summary table of the experiments.

**Table 1:**
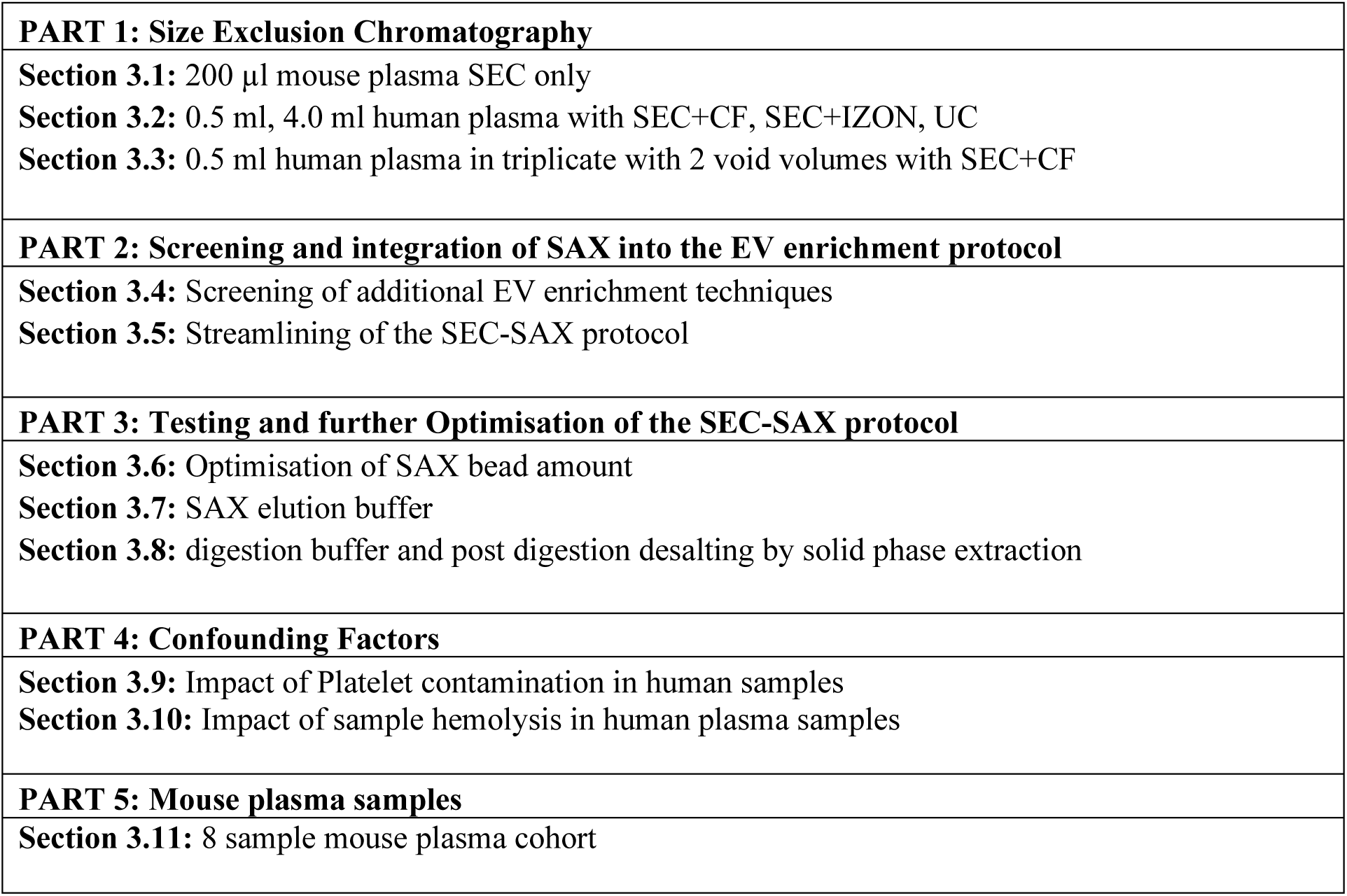
Summary of experiment conducted in the progressive development and testing of the SEC-SAX protocol.

**PART 1: Size Exclusion Chromatography (3.1-3.3)**

### 3.1 Later SEC fractions contain increasing amounts of plasma proteins including albumin

High-abundance soluble plasma proteins obscure the protein signature of EVs during LC-MS based proteomics. Albumin is the most abundant soluble protein in plasma so the reduction of albumin in relation to EVs is important. SEC was performed with 200 µl of mouse plasma using 35nm qEV columns (Izon) to determine the fractions that contained the least albumin and plasma proteins (Figure 1A). Figure 1 indicates that, after removing the void volume, SEC fractions 1-4 contain the least protein content as determined by protein absorbance at 280 nm (Figure 1C), SDS-PAGE (Figure 1B) and albumin as shown via western (Figure 1D). Theoretically these early fractions should contain the extracellular vesicles (EV). However, EV markers could not be detected by western blot likely due to the diluting effect of SEC (data not shown).

**Figure 1:**
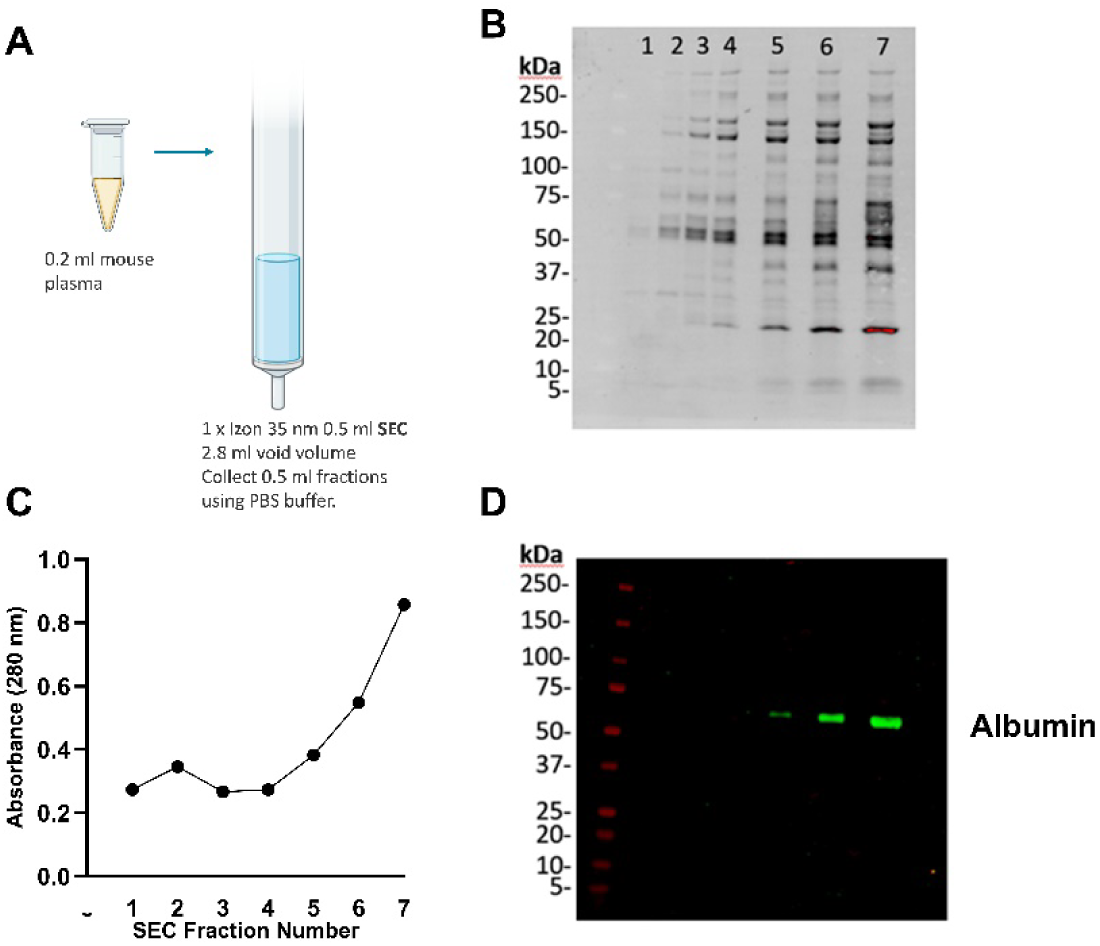
Size Exclusion Chromatography (SEC) of 200 µl Mouse Plasma. Mouse plasma (200 µl) was fractionated using SEC to enrich extracellular vesicles (EVs), as shown in 1A. Fractions 1 through 7 were collected and analyzed for protein content by SDS-PAGE (1B) and by measuring absorbance at 280 nm (1C). Western blot analysis for albumin (1D) demonstrated that albumin was undetectable in fractions 1-4.

### 3.2 SEC + CF allow for detection of CD63 and Syntenin-1 via western blot and partially satisfies MISEV 2018 minimal information for EV studies

As EV markers were below the detection limit by western blot in individual SEC fractions isolated from 200 µl of mouse plasma, we increased the starting plasma volume to 4 ml and included a concentration step (Veerman et al., 2021). Human plasma was used due to the limited amount of plasma available from a single mouse. The human plasma was processed in 0.5 ml lots as this was the maximum recommended input amount for the 35nm qEVoriginal columns (Izon). Plasma was processed using the Automatic Fraction collector (AFC) using a 2.8 ml void volume. Fractions 1-4 were pooled and concentrated from a total of 4 ml of plasma (8 x 0.5 ml lots of plasma) using a single 100,000 Da molecular weight cut off (100kDa MWCO) centrifugal filter, to achieve a final volume of 250 µl as shown in Figure 2A.

**Figure 2:**
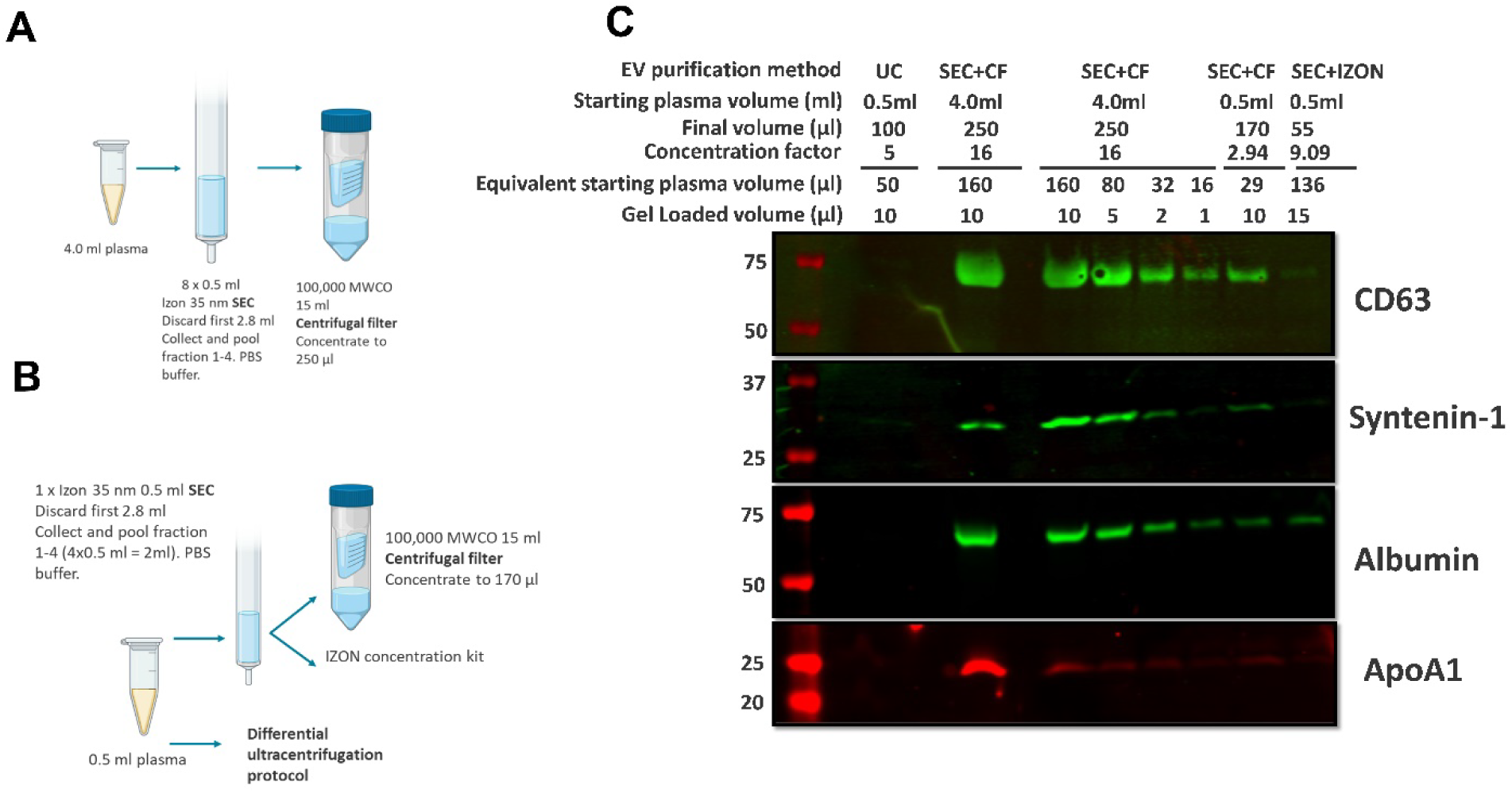
Workflow for EV enrichment from 4 ml (A) and 0.5 ml (B). EV markers (CD63 and Syntenin-1) and plasma contaminant proteins (Albumin and ApoA1(shown in red)) by western blot analysis (panel C). Starting plasma volume, final volume post concentration with the centrifugal filter and gel loading volume were used to calculate the concentration factor and equivalent plasma starting amounts indicated above figure.

The EV markers CD63 and Syntenin-1 were detected in the concentrated SEC material from 4 ml of plasma (Figure 2C). Decreasing amounts of concentrated SEC material were also analysed by western blot, as the objective is EV enrichment from small volumes of plasma. The original starting volume of plasma was calculated as a proportion of the total 4 ml of plasma and the details are shown (Figure 2C). Although a useful guide on the limit of detection, this approach does not take into account sample losses that occur when working with smaller starting volumes so EVs were also purified from 0.5 ml of plasma by SEC plus centrifugal filtration (SEC+CF) (Figure 2B). Comparison with EV purification from 0.5 ml of plasma using ultracentrifugation (UC) and using an alternative concentration method for the SEC material (Izon concentration kit) were also included.

CD63 was difficult to detect and a combination of two CD63 antibodies was used to amplify the signal. As shown in Figure 2C, CD63 was not detected when a starting volume of 0.5ml of plasma was purified using UC, although a faint band for Syntenin-1 was observed. Concentrating 4ml of SEC fractions 1-4 via CF (SEC+CF) resulted in the detection of both EV markers, CD63 and Syntenin-1. However, there was also identifiable bands for albumin and ApoA1 indicating protein contamination. Reducing the volume of SEC+CF material loaded on to the gel (10 µl, 5 µl, 2 µl and 1 µl) resulted in a gradual decrease in the CD63, syntenin-1 and albumin. All proteins were still detectable at all volumes, while ApoA1 was almost undetectable when loading 2 µl and 1 µl.

A scale down version of the SEC+CF protocol was performed using a single 0.5 ml aliquot of plasma (Figure 2B). Loading 10 µl of the sample resulted in detectable levels of the EV markers CD63 and syntenin-1, along with albumin and faint traces of ApoA1. EVs were also purified from a single aliquot of 0.5 ml of plasma by SEC and concentrated using the Izon concentration kit as a potential alternative to the centrifugal filter. Very faint bands for both CD63 and syntenin-1 were observed. There was a detectable band for albumin. Almost no ApoA1 was present although the ApoA1 signal was generally weak for all samples.

SEC + CF protocol allowed for detection of CD63 and Syntenin-1, proteins that fall into category 1 (GPI-anchored membrane associated proteins) and category 2 (cytosolic proteins recovered in EVs), respectively, as defined by the MISEV2018 guidelines for minimal information in EV studies.). Albumin and lipoprotein ApoA1 are major EV contaminants and were measured as a purity control. Albumin was clearly visible, whereas ApoA1 appeared only as faint bands. This demonstrates that SEC purification with concentration by centrifugal filtration of EVs from plasma at least partially satisfy the requirements for purification of EVs from plasma.

### 3.3 EVs purified from plasma by manual SEC +CF satisfy the MISEV 2018 minimal information for EV studies: SEC void optimisation in triplicate

During the development of this protocol the manufacturer changed the recommended void volume (the initial buffer volume discarded prior to collecting EV-containing fractions) from 2.8ml to 2.7ml. We investigated the potential effect of void volume by manual SEC in triplicate. Figure 3A outlines the protocol and following downstream analyses. The effect of changing void volume by 100 µl (2.8 ml versus 2.7 ml) resulted in differences in protein concentration only after fraction 5 (Figure 3B) based on absorbance at 280nm. Therefore, fractions 1-4 continued to be collected, pooled and concentrated as the EV containing fractions and used for further analyses. The absorbance at 280nm was compared in this current experiment (Section 3.3) with manual SEC (2.8 ml void) with the previous samples prepared with the Automatic Fraction Collector (AFC) (Section 3.2) which utilised the same batch of plasma. The protein content, measured by A280nm, from SEC fraction 3 onwards was significantly higher and more variable when using the AFC compared to manual collection (Figure 3C). From this point onwards all SEC was performed manually using analytical pipettes to minimise variation in the sample preparation.

**Figure 3.**
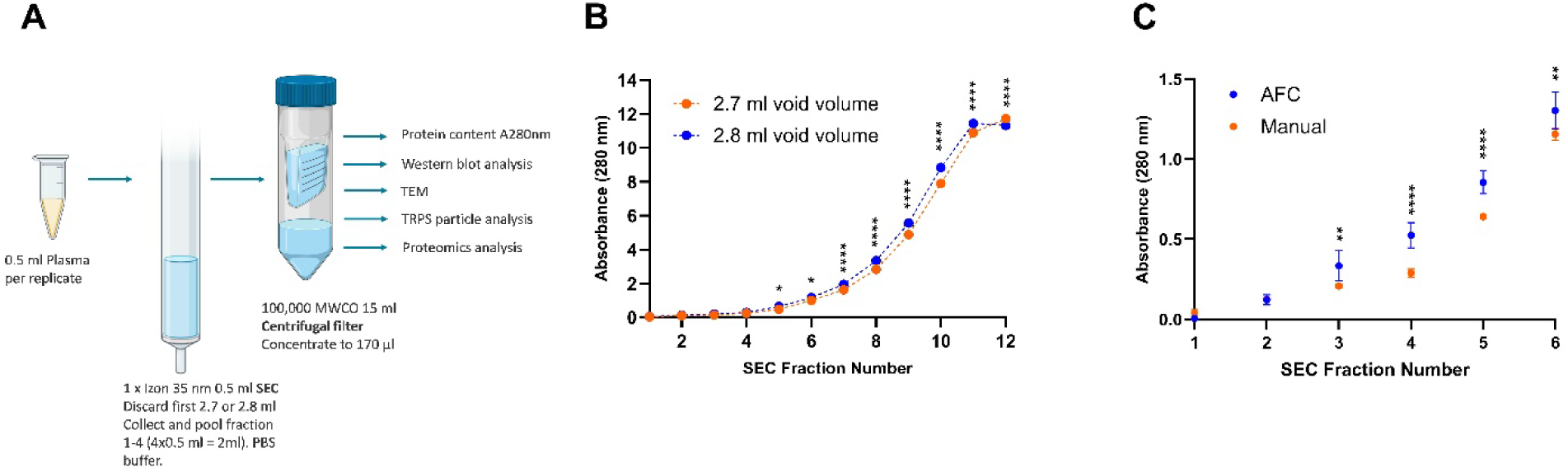
Workflow for optimization of void volume (A), determining protein content of SEC fractions after manual pipetting with 2.7 or 2.8 ml void volume by measuring absorbance at A280 nm (B) or using the automatic fraction collector with a 2.8 ml void volume (C). Data is presented as Mean and SD n=3 for each condition. Error bars (SD) are smaller than symbol for manual pipetting. Statistical significance calculated using a 2-way ANOVA with Mixed-effects model and Tukey correction for multiple comparisons (B,C). Assumption of normality was passed for (C) but not (B). P-value <0.05 (*), <0.01 (**), <0.001 (***), <0.0001 (****).

In Figure 4A, western blot analysis indicated that the plasma protein albumin is more abundant in the EVs purified using the 2.8 ml void volume (Fraction 1-4b), compared with those purified using a 2.7 ml void volume (Fraction 1-4a). This observation remained when comparing fractions 5b, 6b and 7b, with fractions 5a, 6a and 7a, respectively. The positive control (+), unpurified plasma (0.5 µl), was included to show the large amount of albumin contained in the original plasma sample. This demonstrates that SEC removes a large proportion of albumin.

**Figure 4:**
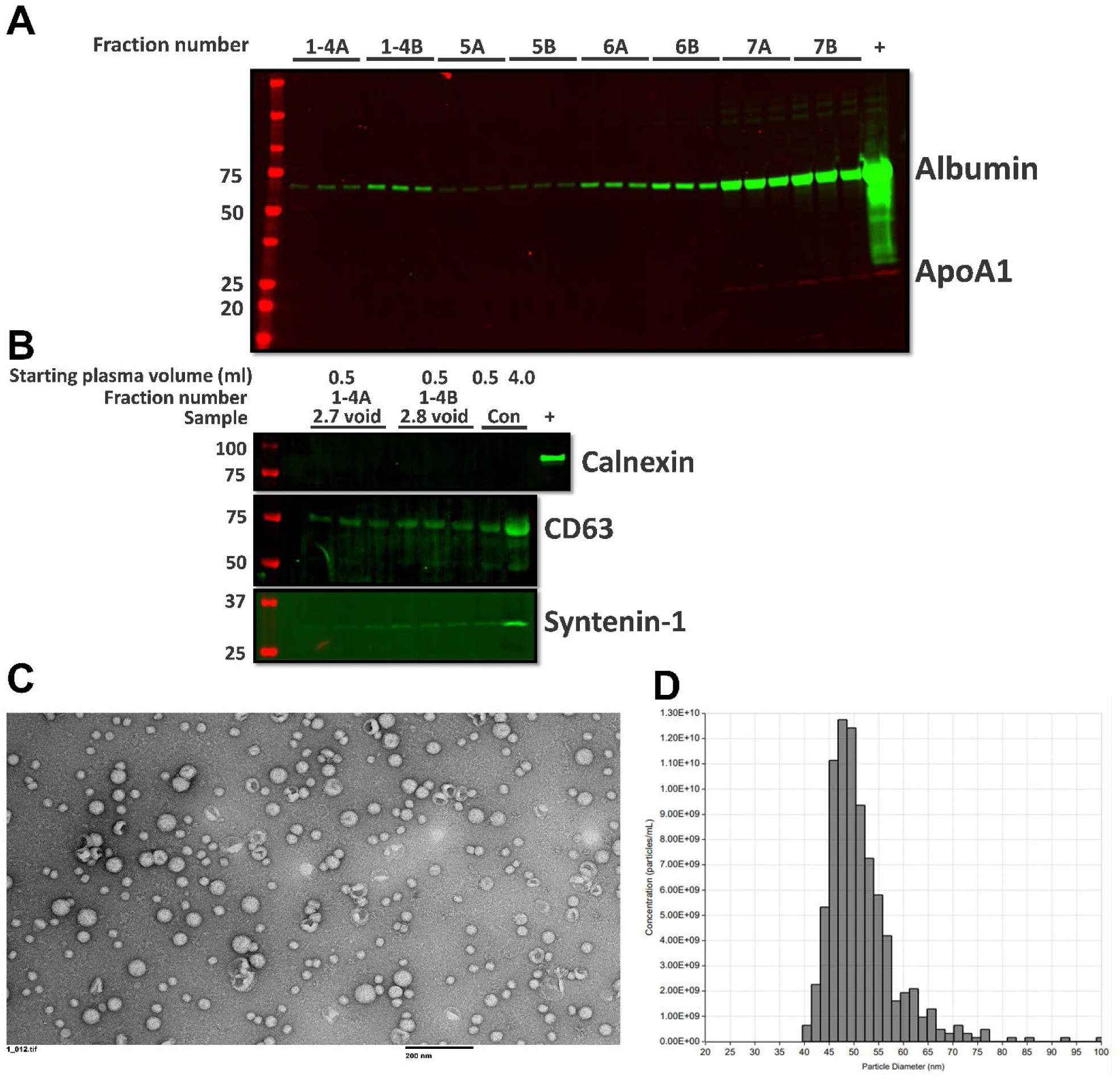
SEC Void optimisation. Pooled SEC fractions 1-4 collected using a 2.7 mL void volume (1-4A) and a 2.8 mL void volume (1-4B) were prepared in triplicate from 0.5 mL plasma. Western blot analysis of contaminant proteins Albumin (green) and lipoprotein ApoA1 (red) (A) with 0.5 µl plasma included as a positive control (+). EV markers CD63 and syntenin-1 and small EV exclusion marker Calnexin (B). Samples (0.5 ml and 0.4 ml) from Experiment 3.2 included as a western blot control and 2µl platelet lysate included as a calnexin positive control (+). Electron microscopy (C) visualisation and size and concentration histogram (D) of EV representative SEC sample (Void 2.7 ml).

The EV markers Syntenin and CD63 (Figure 4B) were detected in all three samples processed with either a 2.7 ml or 2.8 ml void volume. The bands for the EV markers appear fainter and potentially less consistent in the 2.7 samples compared with the 2.8 ml samples. Calnexin was not detected in any sample. Platelets (+) were included as positive control for the calnexin western blot. Calnexin is an exclusion marker for small EVS should not be detected to satisfy the MISEV2018 (Thery et al., 2018). Regardless of the void volume, the SEC+CF isolation of EVs did not contain calnexin, supporting the isolation of small EVs. To further satisfy MISEV2018 guidelines we determined the shape and size of the isolated EVs using electron microscopy (Figure 4C) as well as the concentration and size of the EVs using Tunable Resistive Pulse Sensing (TRPS) (Figure D) (Thery et al., 2018). Both data sets in Figure 4C and 4D are from samples processed with a 2.7 ml void volume. The size by TRPS is consistent with small EVs and the electron micrograph shows particles with typical cup-shaped morphology.

### Proteomics analysis of EVs purified from plasma by manual SEC and concentrated by centrifugal filtration

After satisfying the MISEV2018 conditions with our manual SEC+CF protocol we next established conditions for LC-MS based proteomics to determine the protein signature of the EVs samples isolated using a 2.7 ml and 2.8 ml void volume, as well as from unpurified starting plasma (Supplemental Data S1). Figure 5 shows the workflow used to achieve this.

**Figure 5.**
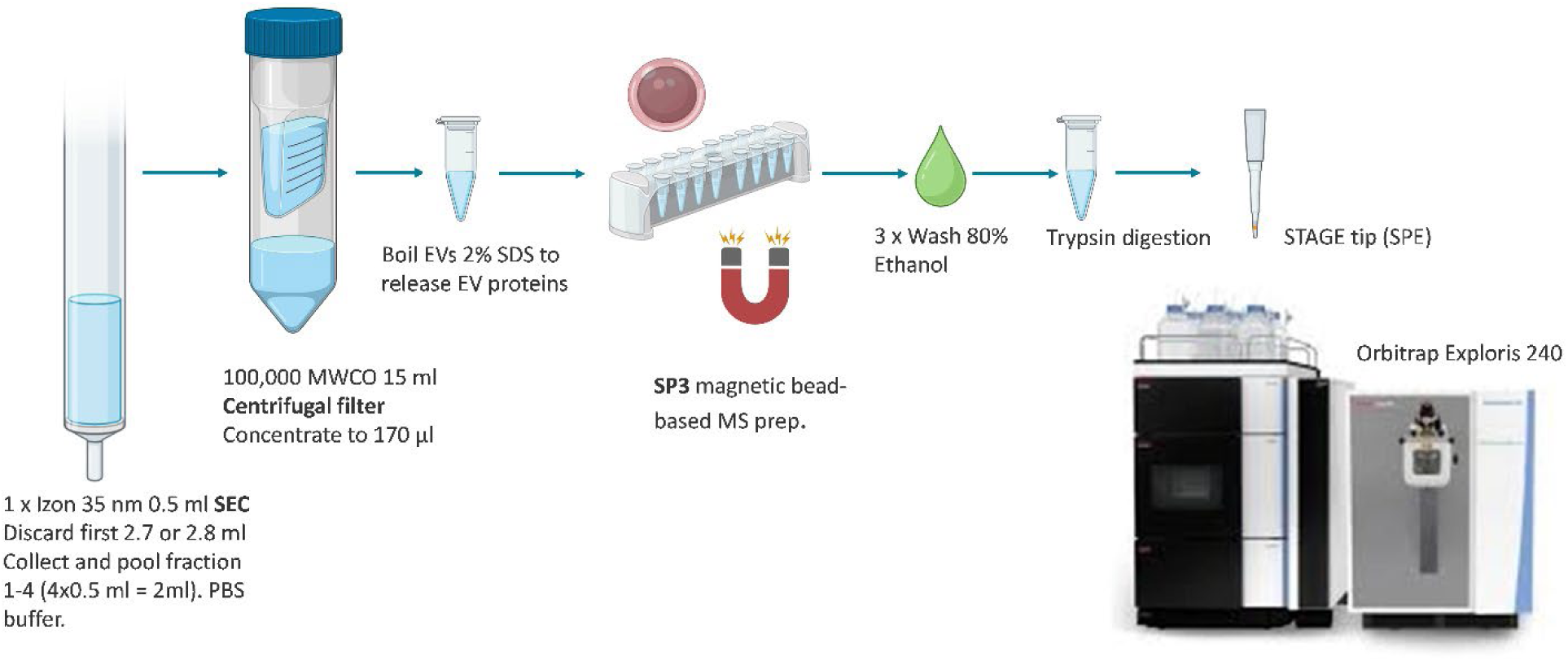
Workflow used to analyse the EVs purified from plasma by manual SEC +CF via LC-MS based proteomics. From the EV samples and starting plasma, 10 µg of material was lysed in 2% SDS for EV lysis/protein solubilisation, this was followed by SP3 single pot sample preparation to remove the SDS. Trypsin digestion was performed, followed by STAGE tip solid phase extraction to desalt the samples.

There was a trend for increased number of quantified proteins with unpurified plasma having the lowest number and SEC purified plasma a higher number based on grouped abundances (Figure 6A). Two different lists were used to identify how many of the quantified proteins were EV markers (STRING analysis (GO:1903561 extracellular vesicle cellular component)(Szklarczyk et al., 2023) and comparison with the Choi list of “Real EV” markers (Choi et al., 2020). The trend for number of EV markers by STRING analysis followed the number of quantified proteins. The number of “real EV” markers was lower in the unpurified plasma. The heatmap (Figure 6B) material shows clear patterns of enrichment and depletion with SEC compared with the starting plasma. The EV markers included Syntenin-1, CD9 and CD81, none of which were identified in the unpurified plasma samples. The EV marker CD63 was not identified in any of the samples despite detection by western blot. All of the 28 EV markers quantified in plasma were also quantified in the SEC purified material indicating that the plasma EV markers are a subsect of the SEC purified material. Additionally, all of the 43 EV markers quantified in the 2.7 ml void SEC samples were also quantified in the 2.8 ml void SEC.

**Figure 6:**
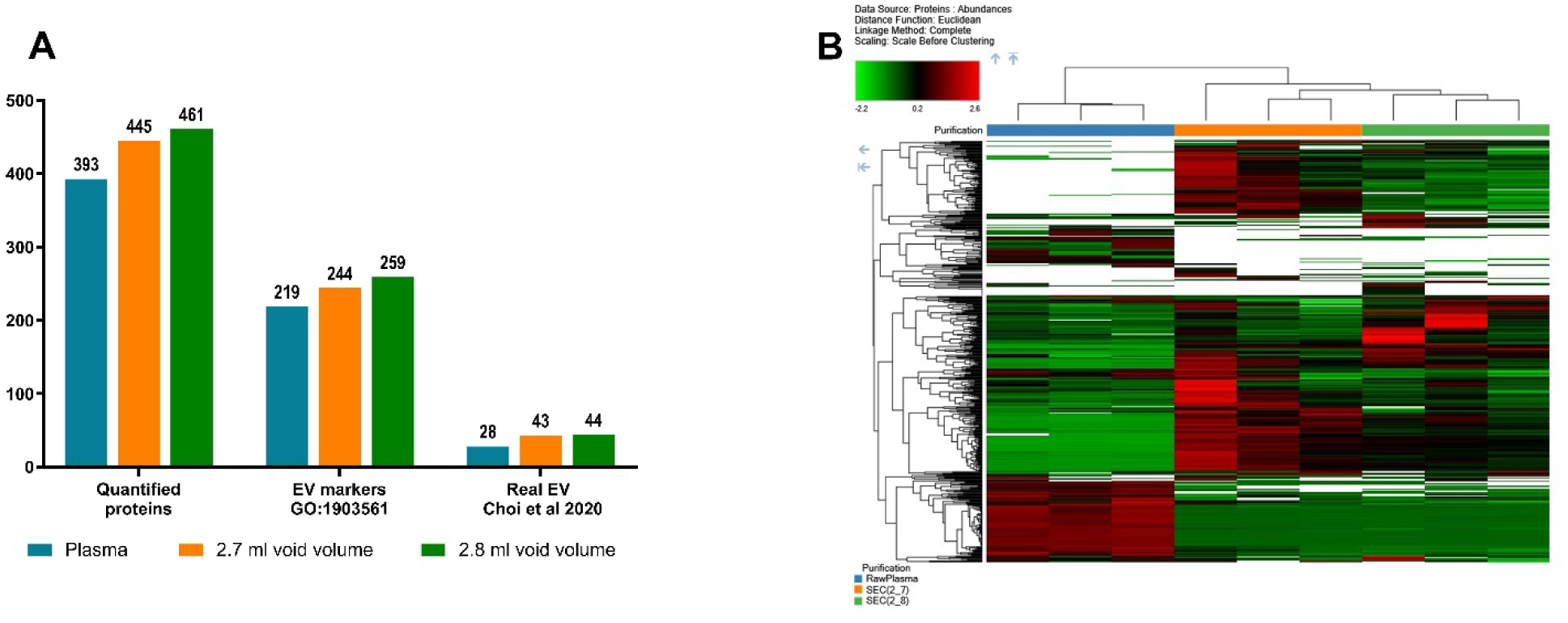
Number of quantified proteins and number of quantified EV markers by STRING analysis and Choi Real EV marker (A). Heatmap comparing plasma and SEC purified plasma (B).

The fold change for albumin, and ApoB were used as an indication of the relative abundance of these proteins (Figure 7). This was used to demonstrate the depletion/enrichment of the plasma contaminants when using SEC+CF compared with the unpurified plasma, as required in the MISEV2018. The fold change of additional key plasma proteins was used to investigate the impact of SEC purification of plasma. Albumin (ALB) and IgG (heavy chain constant region IGHG2, IGHG3, IGHG4) are representative of soluble proteins and were depleted with a fold reduction ranging from 53 to 173-fold. ApoB lipoprotein, IgM immunoglobulin, hemoglobins (HBA, HBB, HBD) and platelet factor 4 (PF4V1) were enriched ranging from 3 to 22-fold by SEC purification. ApoB is the most abundant protein in the samples and is known to be enriched due to the similar size of VLDL (which is composed of ApoB) and EVs (Sódar et al., 2016; Veerman et al., 2021). IgM immunoglobulin is a very large immunoglobulin pentamer with a molecular weight of ∼a million Daltons and is enriched based on size (Ku et al., 2021; Pan et al., 2021; V. Yin et al., 2024b). The hemoglobins are found in red blood cells (Korff et al., 2025; Peter Klinken, 2002) and PF4V1 is a platelet marker (Burkhart et al., 2012) indicating that SEC enriches for residual cellular components not removed by centrifugation during preparation of plasma.

**Figure 7:**
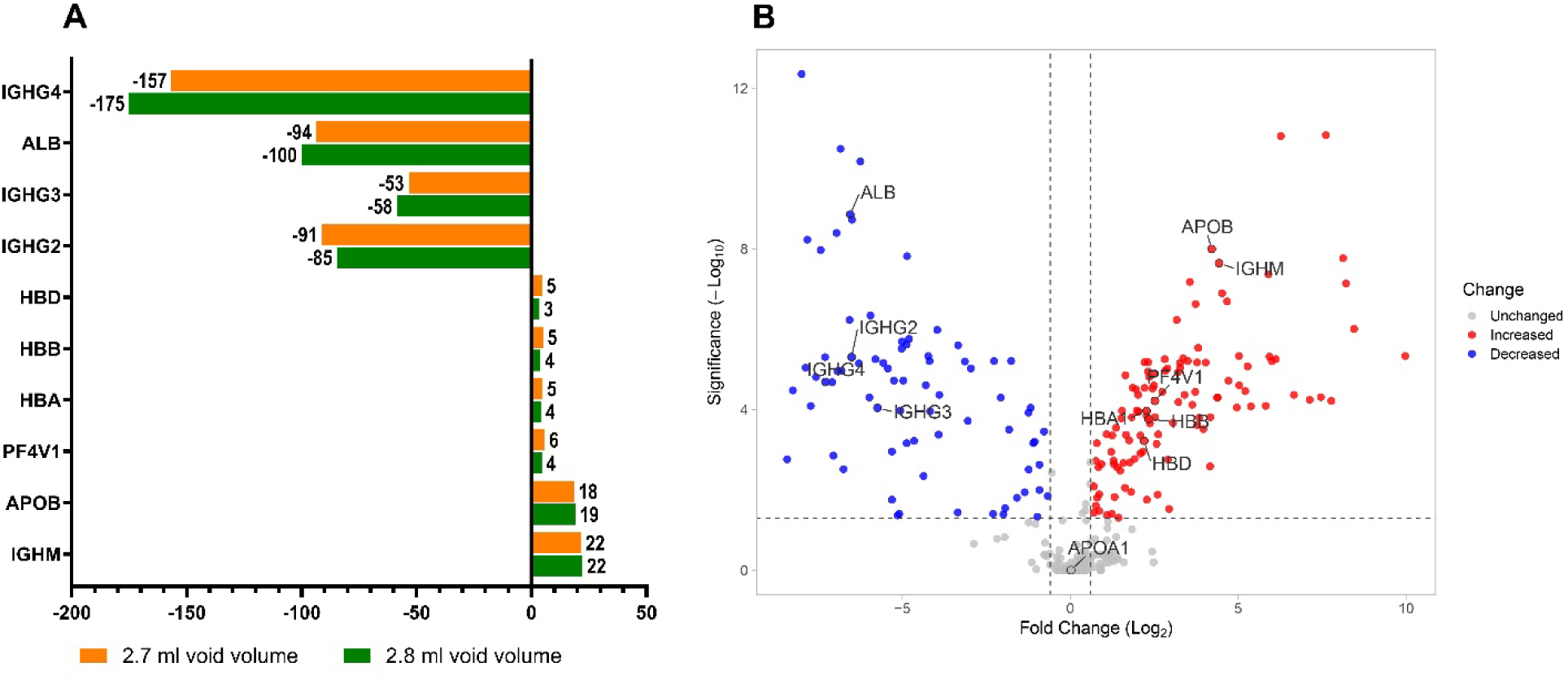
Fold change in plasma contaminant proteins: IgG (heavy chain constant region IGHG2, IGHG3, IGHG4), and IgM immunoglobulins, Albumin (ALB), ApoB lipoprotein and hemoglobins (HBA, HBB, HBD) and platelet marker (PF4V1) for SEC+CF purified EVs compared with plasma via LC-MS (A). Representative volcano plot for SEC (2.7 ml void) purified plasma compared with plasma (B).

The individual abundances were used to calculate intrasample ratios of ApoB and Albumin to EV markers Syntenin, CD9 and CD81 as an indication of EV purity independent of protein concentration (Figure 8). EV purity is defined as the “ratio between EVs and non-EV components” (Coumans et al., 2017). For the intrasample ratios the higher value indicates greater relative contamination, while lower ratios indicate better purity. The EV purity ratios indicate that ApoB has a relative abundance ranging from 9,847 to 53,730-fold for ApoB (Syntenin 9,847 - fold, CD9 23,179-fold and CD81 41,814 - fold) for the 2.7 ml void samples. The corresponding values for the 2.8 ml void samples trended higher, indicating worse purity, with statistically significant differences for all three EV markers (Figure 8A). The EV purity ratios for Albumin range 185 to 904-fold for albumin in relation to the EV markers (Syntenin 185- fold, CD9 435-fold and CD81 783- fold) for the 2.7 ml void samples. These values are much lower than for ApoB, which is consistent with the depletion of albumin by SEC as shown by both Western blot and LC-MS proteomics (Figure 4A). Similarly to the ApoB purity ratios, the values for the 2.8 ml void are trending higher compared with the 2.7 ml void, with the difference reaching significance for syntenin (Figure 8B). Although the differences between the 2.7 and 2.8 ml voids are numerically small, they are significant for the EV purity ratio for Syntenin in relation to albumin and all three EV markers in relation to ApoB. These findings support the idea that intrasample protein abundance ratios can be used as a sensitive indication of EV purity, which are independent of protein concentration. They also demonstrate that void volume directly affects EV purity, underscoring the need for precise and consistent control of this parameter. Taken together, these results indicate that SEC alone does not sufficiently remove ApoB for robust EV proteomic analysis, and that incorporating an additional purification step may be beneficial.

**Figure 8.**
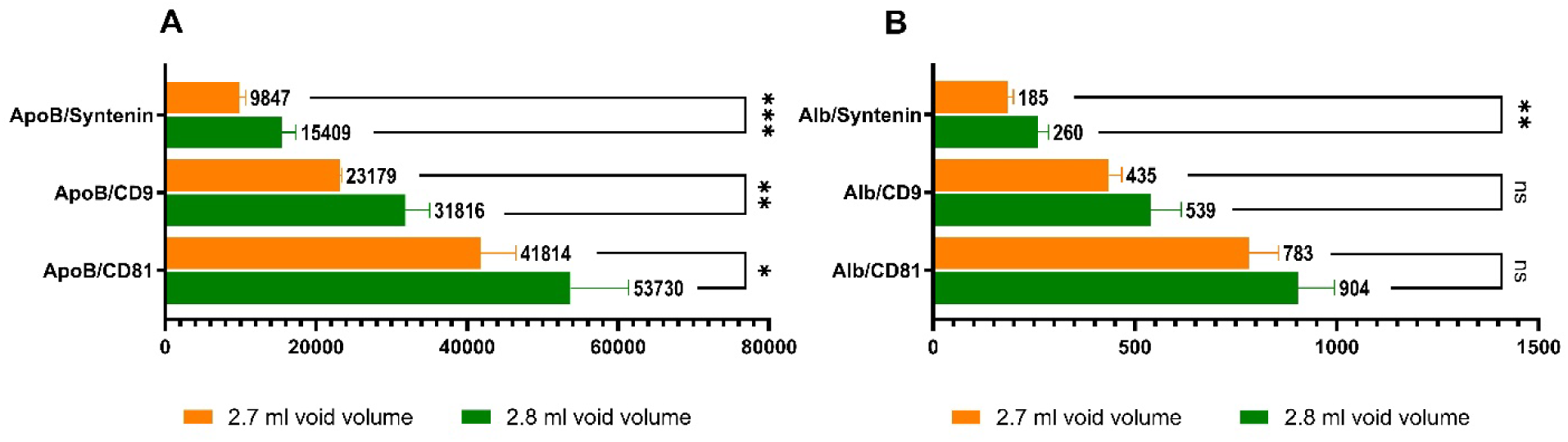
Intrasample EV purity ratios for EV marker proteins Syntenin, CD9 and CD81 in relation to plasma contaminant proteins Albumin (Alb) and Apolipoprotein B (ApoB). Statistical significance calculated by 2-way ANOVA with Holm-Šídák’s multiple comparisons test from Log2 transformed ratios. Data presented as Mean and SD of non-transformed data. P-value <0.05 (*), <0.01 (**), <0.001 (***), <0.0001 (****).

**PART 2: Screening and integration of SAX into the EV enrichment protocol (3.4-3.5)**

The two experiments in Part 2 are screening experiments so were not conducted in triplicate and statistical tests were not used.

### 3.4 Enrichment screen for additional EV enrichment techniques

A pool of concentrated SEC material (SEC+CF) was used as the starting point to test the added effectiveness of three EV enrichment techniques (Figure 9): Anion Exchange magnetic beads (SAX), CD9, CD63, CD81 magnetic beads (immune affinity capture, IAC) and the EV Concentration kit (IZON). Additionally, the SAX beads were tested with precleared plasma without SEC+CF. All samples were tested in duplicate with the exception of the IZON concentration kit (Supplemental Data S2). The workflow for the EV enrichment is shown in Figure 9A. The IZON concentration kit was only included for completeness after being found in Section 3.2 (Figure 2) to show reduced EV enrichment by western blot for EV markers CD63 and Syntenin compared with SEC+CF.

**Figure 9:**
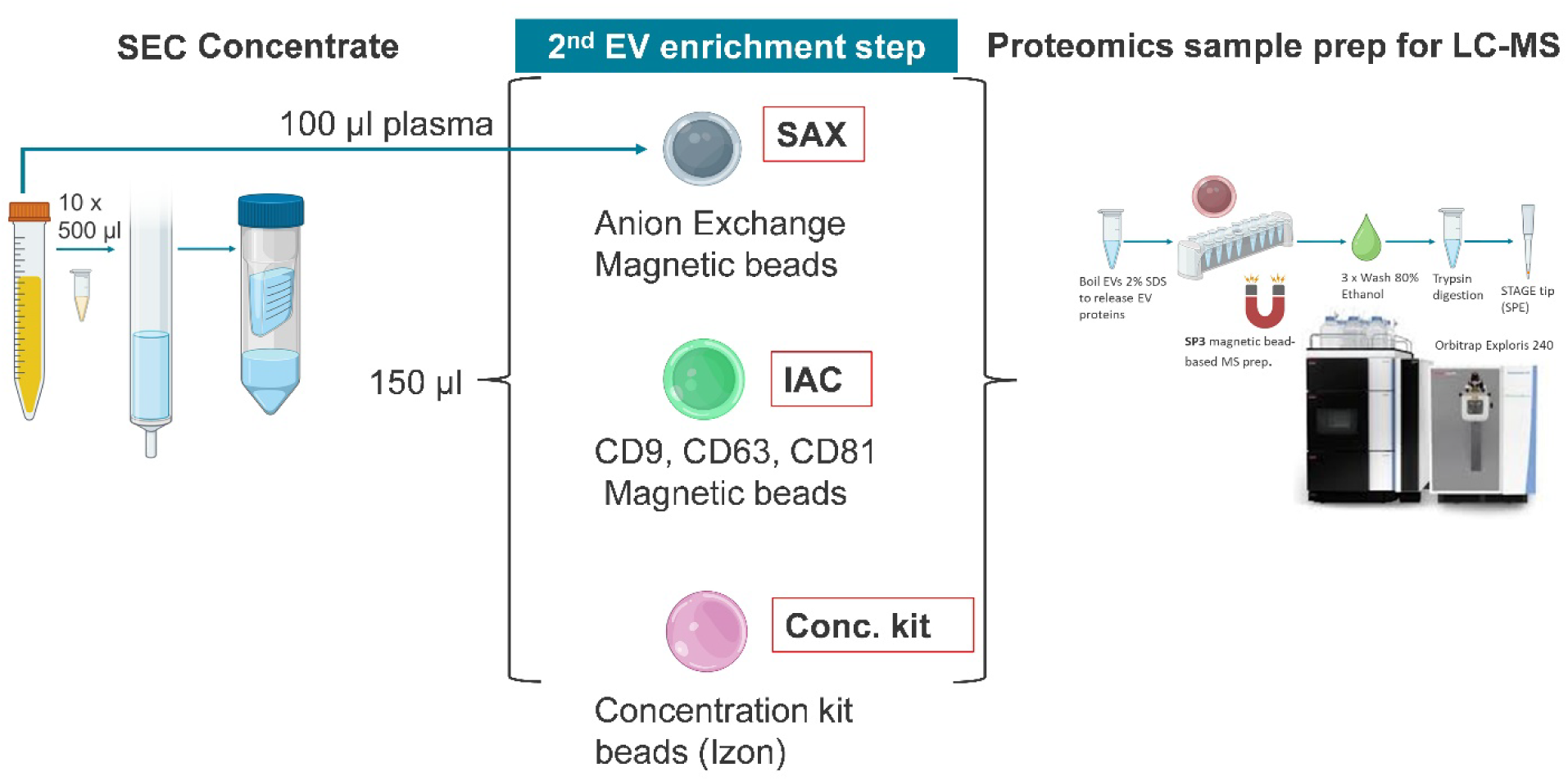
Workflow for screening additional EV enrichment techniques

The number of EV markers identified per sample as defined by the “real EV” marker list was used as the primary readout of EV enrichment. The number of EV markers is plotted in Figure 10A and is further separated into high and low abundance proteins to identify the best EV enrichment technique. Combining SEC with SAX (SEC-SAX) more than doubled the total number of EV markers detected from 37 to 92 compared with SEC+ CF alone. The majority (88 of 92) of the EV markers detected with SEC-SAX were high abundance EV markers, supporting SEC-SAX as the most effective method and justifying its selection for further optimisation. None of the other enrichment techniques tested yielded comparable increases in total EV marker numbers compared with SEC + CF alone. Both SAX alone (38 markers) and SEC-IAC (33 markers) showed evidence of EV enrichment, as indicated by an increase in high-abundance EV markers compared with SEC alone (12 markers). The presence of the key EV markers CD9, CD81 and syntenin was used as a secondary method to confirm EV enrichment (Figure 10C). All three markers were confirmed as high abundance proteins for the combination of SAX with SEC confirming its suitability of this combined method as an EV enrichment technique. The categorisation of proteins as high or low abundance is based on peak intensity. As such, the results were confirmed with the actual quantified proteins using protein abundances (Figure 10B). The SEC-SAX method had the highest number of quantified proteins (522 proteins), 54% of which were identified as EV markers by STRING analysis (299 proteins) and 71 were “real EV” markers.

**Figure 10:**
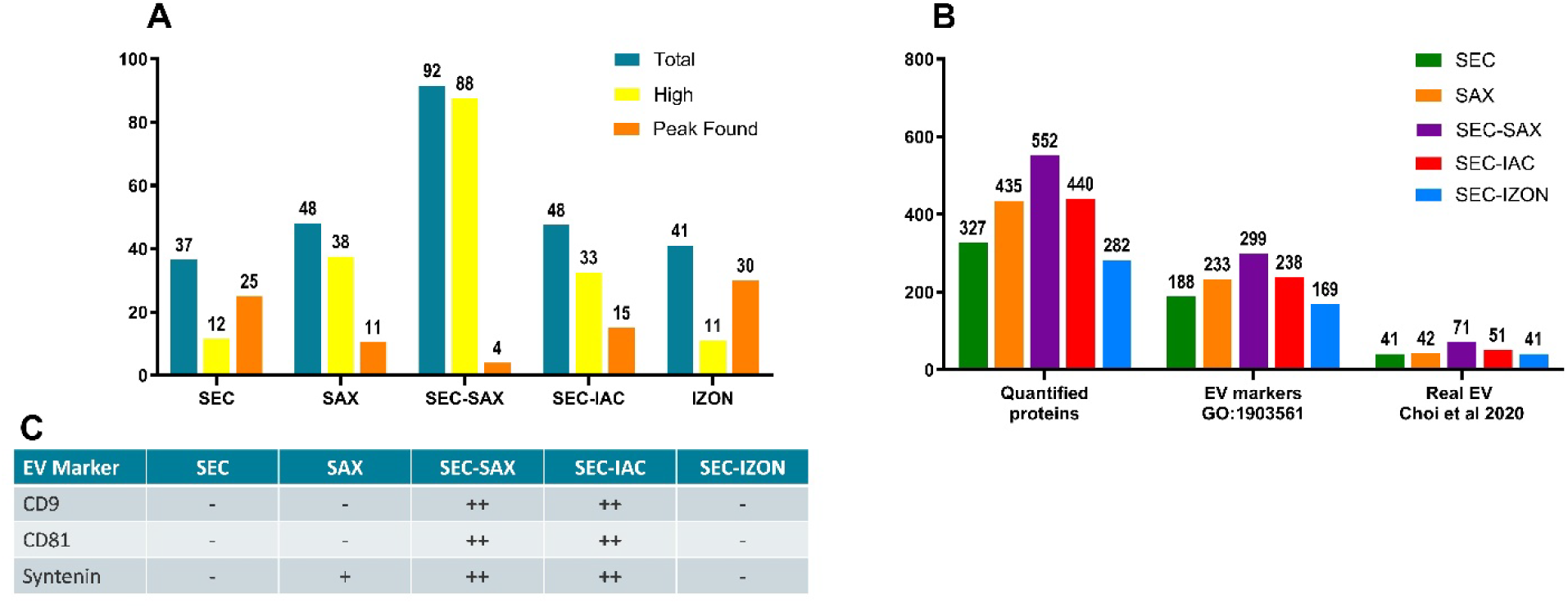
Comparison of additional EV enrichment techniques. “Real” EV markers per enrichment technique (average of duplicate samples) categorised by abundance based on peak intensity (A). Quantified proteins, EV markers (STRING) and “Real” EV markers based on grouped abundances (B). Abundance of EV markers Syntenin-1, CD9 and CD81 based on peak intensity (- not detected, + peak found, ++ high abundance) (C).

A heatmap of all the samples (Figure 11A) shows clear enrichment by SEC-SAX compared to SEC alone with enriched proteins shown in red and relatively depleted proteins in green. The heatmap also indicates enrichment by SEC-IAC and the SAX beads with plasma used as starting material although neither is as consistent as the SEC-SAX. A PCA plot (Figure 11B) shows that SEC-SAX is clearly different to the starting SEC material and the other methods. There is no additional enrichment benefit under the conditions tested for the Izon concentration kit based on the lack of difference from the starting SEC material in the PCA plot and heatmap.

**Figure 11:**
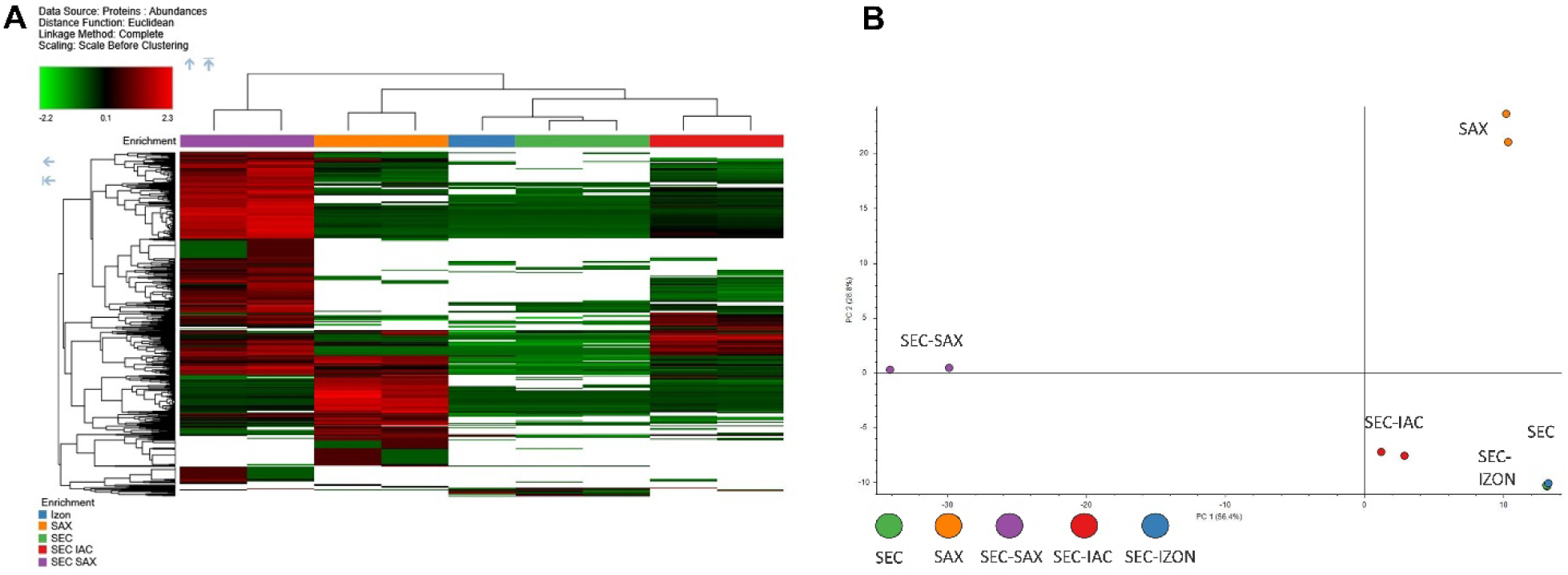
Heatmap (A) and PCA plot (B) comparison of different enrichment techniques.

The combination of SEC with CD9, CD81, CD63 beads (SEC+IAC) appears to be the second most effective enrichment method. It shows enrichment of key EV markers (Figure 10C) and increased proteome depth, as indicated by the number of quantitated proteins (Figure 10B) of which 54% (238 of 440) were EV markers by STRING analysis and 51 were “real EV” markers. In contrast, enrichment using the SAX beads alone is less effective for EV enrichment, although it resulted in greater proteome depth compared with SEC alone (Figure 10B). The key EV markers were not quantified in SAX only enrichment and only syntenin was identified as a low abundance protein (Figure 10C).

### 3.5 Streamlining the SAX-SEC protocol

#### Centrifugal filtration is unnecessary in the SEC-SAX protocol when Bis-Tris propane buffer is used as the SEC buffer

The number of EV markers quantified in each sample based on the “Real EV” marker list (Choi et al 2020) in addition to the quantification of EV markers Syntenin, CD9 and CD81 was used to assess the impact of removing potentially redundant protocol steps (Supplemental Data S3). Figure 12 shows the workflow variations tested.

**Figure 12:**
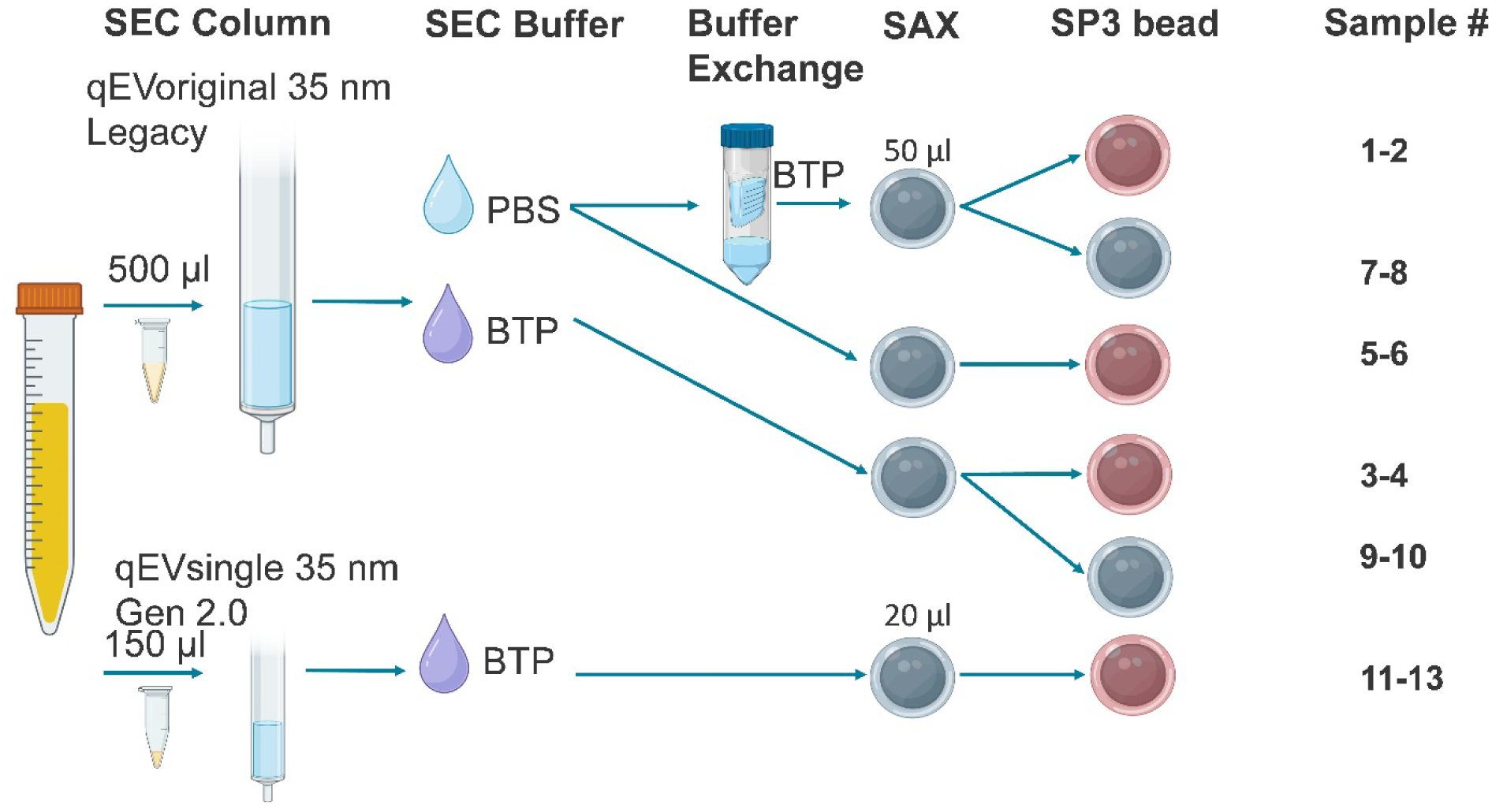
Workflow of the different combinations of buffers (PBS or BTP), SP3 beads (SAX beads (grey) or Cytiva carboxylate beads (brown)), SEC column scale (500 µl or 150 µl), SAX beads amount (20 µl or 50 µl) and buffer exchange by centrifugal filter tested to remove redundant protocol steps.

Samples 1 and 2 use the same method established in section 3.4 and act as the standard for this experiment. The objective was to remove redundant steps, specifically if the centrifugal filtration concentration/buffer exchange step could be removed by substituting the SEC buffer with Bis-Tris Propane and the results are shown in Figure 13. This was tested with samples 3 and 4 and the number of quantified EV proteins is maintained at the same level as the standard indicating that this substitution in the protocol produces equivalent results. Bis Tris Propane is more expensive than PBS and is not available in a premixed tablet format like PBS so is more complicated to prepare. Samples 5 and 6 test if the centrifugal filter concentration step could be removed while continuing to use PBS as the SEC buffer. The number of EV markers quantified (82 and 85 markers) was less than for the standard method (96 and 96 markers) indicating that this combination is not ideal and the BTP buffer is preferred. Using two different types of beads during the protocol is more expensive and introduces additional handing steps and potential for variation. The SAX beads have been used previously for the SP3 protocol (Wu et al., 2025) so samples 7-10 test their use for both EV enrichment and mass spec. preparation. Samples 7-8 were prepared the same as for the standard samples 1-2 with just the substitution of the SP3 beads with the SAX beads. The number of EV markers quantified was reduced when using the SAX beads (82 and 85 markers) in place of the SP3 beads (96 and 96 markers). This suggests that the SAX beads do not capture the proteins as effectively as the standard SP3 beads and the use of 2 types of beads in the protocol is justified. Samples 9 and 10 were prepared the same as samples 3 and 4 using the BTP buffer for SEC and omitting the centrifugal filter. The number of EV markers when substituting in the SAX beads (95 and 99 markers) for the SP3 beads in this combination was the same as samples 3-4 (97 and 98 markers) and for the standard method samples 1-2 (96 and 96 markers). However, CD81 was not quantified in either sample 9-10 or sample 8 confirming that the use of two types of beads is justified in the protocol. We speculate that the removal of the centrifugal filter with samples 9-10 results in reduced sample losses compared with samples 7-8 which compensates for the reduced effectiveness of the SAX in the proteins capture step of the mass spec prep resulting in better EV markers numbers than for samples 7-8. However, the decreased sample loss is insufficient to compensate for the low abundance CD81 marker. The final 3 samples (11-13) in this experiment test the use of the smaller 150 µl volume SEC columns with the new Gen 2.0 resin (qEVsingle 35nm Gen 2.0). These were tested with the BTP SEC buffer without the centrifugal filter as we speculated that this would be the best combination when planning the experiment. A reduced amount of SAX beads of 20 µl compared with 50 µl was used to account for the reduced amount of input plasma amount of 150 µl compared with 500 µl. Additionally, half the volume of loading solvent was used to resuspend the desalted peptides but with the same injection volume. This was to compensate for the reduced plasma amount by effectively doubling the proportion of peptides injected onto the LC-MS. The only difference between the 3 samples is the number of fractions collected and pooled. The void volume and fraction volume was based on the manufacturers recommendations and volumes equivalent to 4, 5 and 6 fractions were collected (samples 11, 12, 13 respectively). The number of EV markers was highest (109 markers) with the collection of equivalent to 4 fractions collected as a pool which is the same as for the method established with the larger 0.5 ml columns (qEV original 35nm Legacy resin). Thus, minor adjustments to SAX bead amount and peptide injection amount is sufficient to scale down and transfer the method to the smaller columns without loss of EV marker quantification.

**Figure 13:**
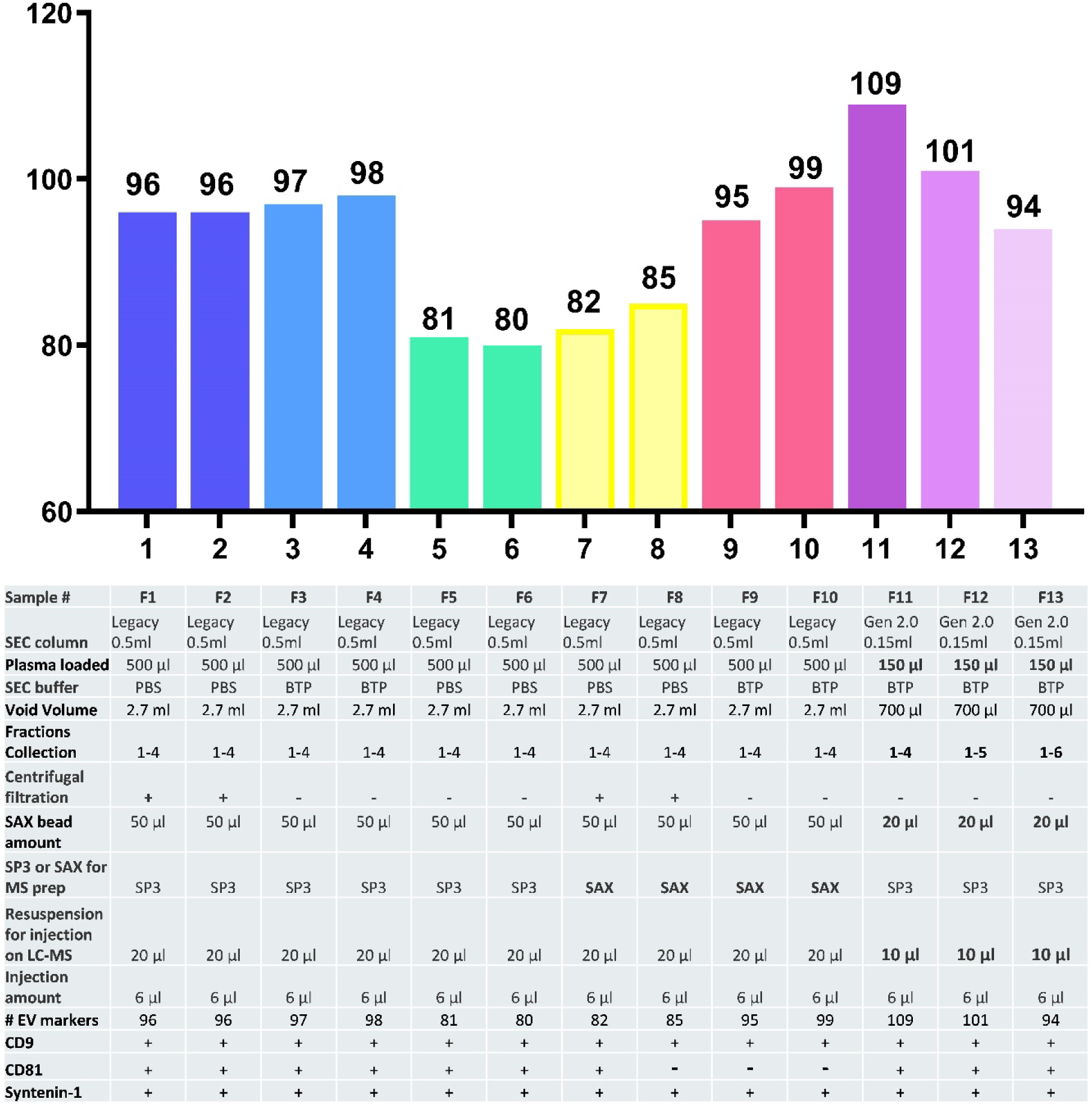
“Real” EV marker number (A) plot and table summarising the variable parameters for each sample (B).

**PART 3: Testing and further Optimisation of the SEC-SAX protocol (3.6-3.8)**

PART 3 focused on testing the SEC-SAX protocol after integration of the SAX magnetic beads into the EV enrichment protocol and removal of redundant steps in PART 2. All samples were analysed in triplicate to allow for a comprehensive analysis for each parameter tested and a diagram of the workflow used for these experiments is shown in Figure 14.

**Figure 14:**
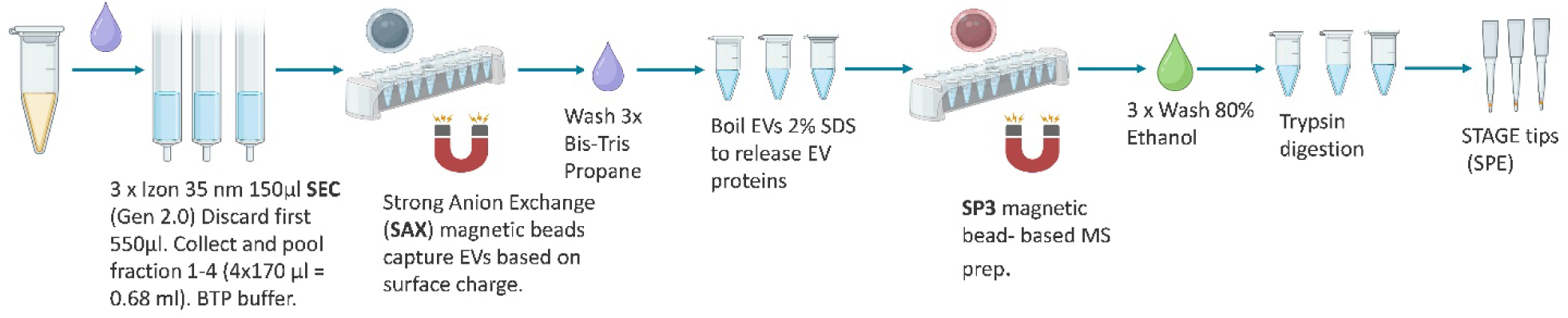
General workflow for the SEC-SAX protocol in triplicate

### 3.6 SAX beads amount optimisation

Two different amounts of SAX beads (20 µl or 40 µl) were tested for EV enrichment from the SEC eluant to determine whether using more SAX beads would capture more EVs, or whether a smaller bead volume could achieve comparable results while reducing costs (Supplemental Data S4). Both bead amounts produced very similar results based on number of quantified proteins and number of EV marker by both STRING analysis and comparison with the “real EV” marker list (Figure 15A).

**Figure 15:**
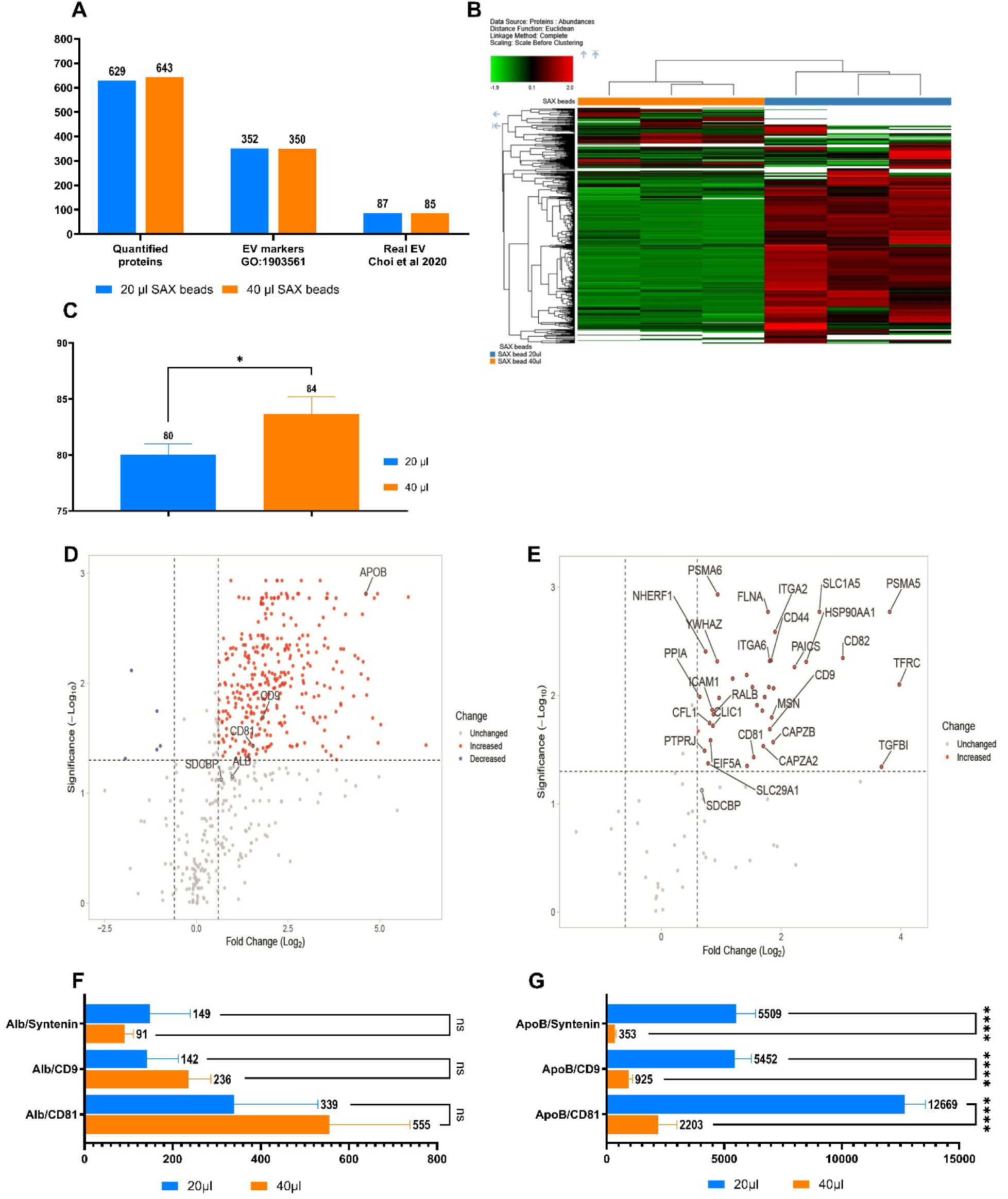
Grouped abundances plot (A), heatmap (B) “real” EV marker number plot (C), volcano plot for all proteins (D), volcano plot for “Real” EV markers (E), and intrasample ratio plots for albumin (F) and Apolipoprotein ApoB (G) for comparison of 20 µl and 40 µl SAX beads. Statistical significance calculated by unpaired T-Test (Plot C), 2-way ANOVA from Log2 transformed ratios (plots F,G) with Holm-Šídák’s multiple comparisons test. Data presented as Mean and SD of non-transformed data. P-value <0.05 (*), <0.01 (**), <0.001 (***), <0.0001 (****).

Further analyses were conducted to determine if a significant difference exists between the groups. The heatmap (Figure 15B) indicates greater protein enrichment (red) with less (20 µl) bead amount. This is confirmed by the volcano plot (Figure 15D) of the both the total proteins and the “real EV” proteins (Figure 15E), with a trend for greater enrichment of all proteins including both contaminant and EV marker proteins. However, when examining samples individually, 40 µl SAX of beads resulted in a significantly greater number of quantified EV markers, (Figure 15C), averaging 84 EV markers compared with 80 for the 20 µl condition. The EV purity ratios for albumin (Figure 15F) and ApoB (Figure 15G) show no significant difference between bead amounts for albumin. However, the EV purity ratio relative to ApoB was significantly improved with 40 µl of beads across all three EV markers. This suggests that SAX beads preferentially enrich for EVs over ApoB and that increasing SAX bead volume results in better EV enrichment. However, the overall yield of proteins recovered is reduced with more SAX beads. Comparison of the EV purity ratios determined for SEC alone (Figure 8A, Section 3.3) shows a large improvement with the addition of SAX beads to the EV protocol in relation to ApoB but not albumin. The 20 µl SAX bead amount was selected for further experiments to prioritise protein yield over EV purity given the equivalent number of EV markers quantified when the data was combined into grouped abundances (Figure 15A). This data shows the strength of the using biological replicates, as pooling data from three replicates substantially increases the number of quantifiable EV markers (compare panels A and C).

### 3.7 SAX bead elution buffers

The buffer used for the SAX bead elution was further investigated as data from the comparison of SAX beads amount (section 3.6) showed reduced protein recovery with increased SAX bead amount. Two salt concentrations (400 mM and 800mM) NaCl were tested for improved elution as salt is often used as elution method for anion exchange. Additionally, the inclusion of the non-ionic detergent Triton X100 was also tested for improved EV elution as we suspected there may be an interaction between the SAX beads and the anionic SDS detergent used for the standard elution (Supplemental Data S5). The grouped abundance data (Figure 16A) shows similar quantified proteins for all elution buffers although samples eluted with buffers containing 400 mM NaCl (701 proteins for 400 mM and 400 mM+ Triton) had fewer quantified proteins compared with the standard buffer (730 proteins) and the high salt (800mM) buffer with Triton (718 proteins). This trend was also seen in the EV markers by STRING analysis. More “real EV” markers were quantified from the high salt + triton buffer (122 proteins) compared with the standard buffer (118 proteins). Analysis at the individual sample level (Figure 16B) confirmed this with significantly more “real EV” markers quantified from the high salt + triton buffer (120 EV markers) compared with the standard buffer (113 EV markers). While the samples eluted with the 400 mM salt buffer both with added Triton (103 EV markers) and without added Triton (105 EV markers) had significantly fewer EV markers compared to the standard buffer (113 EV markers). The EV purity ratios show reduced purity for CD9 in relation to albumin for the 400 mM NaCl containing buffers. The EV purity ratios in relation to ApoB show improvement for the high salt (800 mM NaCl) with Triton. The inclusion of Triton X100 in the SAX elution buffer in combination with high salt impacts ApoB enrichment whereas the inclusion of 400 mM salt has greater impact on albumin. The heatmap (Figure 16K) shows an obvious lower enrichment of proteins (green) for the high salt with Triton buffer compared to the standard buffer and the 400 mM containing buffers. There is greater enrichment by both the 400 mM NaCl containing buffers compared with the standard buffer. The volcano plots of all proteins (Figure 16E and 16G) and EV markers (Figure 16F and 16H) show that the 400 mM containing buffer elute more protein including both EV and contaminants. This is significant for CD81 and albumin for 400 mM NaCI without Triton and Syntenin (SDCBP) and albumin for 400 mM NaCl with Triton but not ApoB. The Volcano plots for high salt buffer with Triton show a significant reduction for ApoB and an increase for CD81 but no change for Albumin or the EV markers CD9 and syntenin (Figure 16I and 16J). This is consistent with the EV purity ratios which are more significant for CD81 and CD9 than syntenin in relation to ApoB and confirms the sensitivity and utility of the EV purity ratios (Figure 16D). The volcano plot for the high salt buffer with Triton also confirms reduced overall protein enrichment consistent with the heatmap (Figure 16K). This reduced enrichment extends to many EV proteins, so we continued with the standard buffer to prioritise protein recovery (Figure 16J). The 400 mM NaCl containing buffers were not used as they increased the recovery of Alb and reduced the EV purity ratio of CD9.

**Figure 16:**
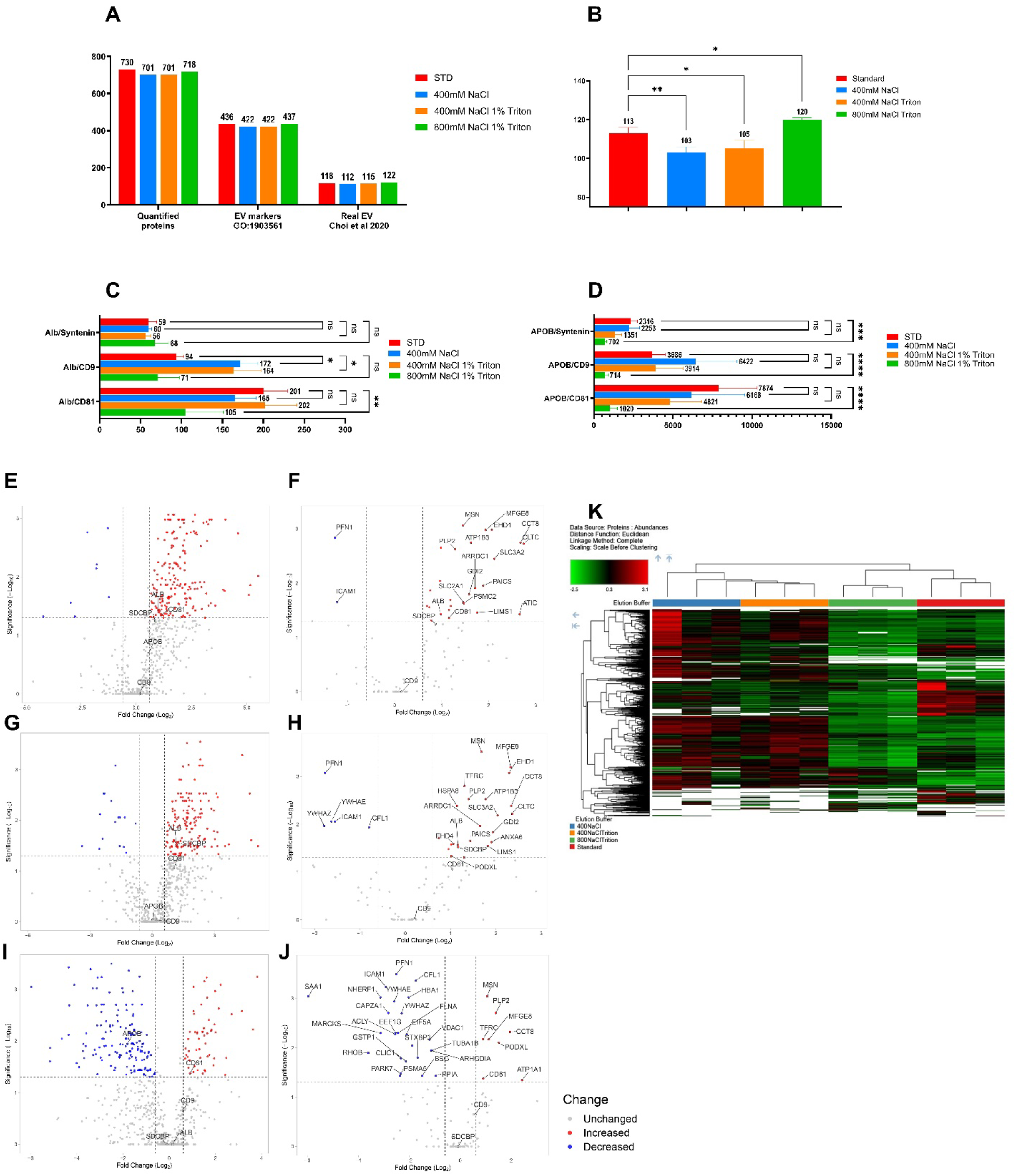
Grouped abundances plot (A), “real” EV marker number plot (B), intrasample ratio plots for albumin (C) and Apolipoprotein ApoB (D) volcano plot for all proteins (E, J,I), volcano plot for “Real” EV markers (F, H, J), and heatmap (K) for comparison of standard elution buffer (100 mM HEPES + 2% SDS) with added 400mM (Volcano E, F) or 400 mM NaCI +1% Triton X-100 (Volcano G,H) and 800 mM NaCl 1% Triton X-100 (Volcano I,J). Statistical significance was calculated by one-way ANOVA (plot B), 2-way ANOVA with Holm-Šídák’s multiple comparisons test from Log2 transformed ratios (plots C,D). Data presented as Mean and SD of non-transformed data. P-value <0.05 (*), <0.01 (**), <0.001 (***), <0.0001 (****).

### 3.8 Trypsin digestion and post digestion clean up

The final step in the SEC-SAX protocol is desalting of the peptides post-trypsin digestion using STAGE tips for solid phase extraction. This experiment was designed to test if the STAGE tip solid phase extraction could be substituted with a technique that required less manual handling. We also tested if the trypsin digestion time could be reduced to just 6 hours. The STAGEtip solid phase extraction step is used to clean up the trypsin digestion and remove the non-volatile HEPES buffer. We reasoned that if the volatile ammonium bicarbonate buffer ±TFE gave the same depth of sequence as the HEPES digestion buffer, the STAGEtip SPE step could potentially be omitted, and the digestion could be dried using a CentriVap to remove the digestion buffer prior to resuspension in the LC-MS loading buffer (Supplemental Data S6). However, the results shown in Figure 17 indicate that the STAGEtips, whether eluted directly into a micro vial insert (STAGEvial) or transferred into the micro vial insert after resuspension in LC-MS loading solvent (STAGEtip), is necessary for best results. Digestion in ammonium bicarbonate with or without TFE (AM or AMT) for 6 or 18 hours resulted in reduced numbers of quantified proteins and EV markers identified by STRING analysis and “Real EV” marker list (Figure 17A). This reduction was even more obvious at the individual sample level (Figure 17B). The standard method quantified 85 “real EV” proteins, compared with 66 (6 h) and 64 (18 h) in ammonium bicarbonate alone, and 53 (6 h) and 61 (18 h) in ammonium bicarbonate + TFE. The heatmap (Figure 17C) indicates that 18-hour digestion in ammonium bicarbonate without STAGEtip desalting provides greater enrichment for some proteins (red signal in the upper half of the heatmap). However, this comes at the expense of presumably low-abundance proteins, which are recovered only when STAGEtip desalting is used (visible in the lower third of the heatmap). This suggests that although STAGEtips may introduce greater sample loss than evaporation-based cleanup, the gain in depth of coverage, due to more effective contaminant removal by SPE, outweighs this disadvantage. Protein abundances plots (Figure 17D) confirmed the lower overall abundance of samples desalted by SPE consistent with higher sample losses but also greater sensitivity. There are also obvious effects of the digestion buffer itself (ammonium bicarbonate with or without TFE) compared with the standard digest buffer of HEPES + TFE but these were not investigated further as the objective was to determine if the STAGEtips were a necessary part of the protocol. From this experiment we concluded that the STAGEtip desalting remains necessary. Importantly, STAGEtips can be eluted directly into the LC-MS micro vial inserts without impact on the EV markers. This represents a minor but useful refinement to the protocol, avoiding the need to transfer peptides between tubes after elution.

**Figure 17:**
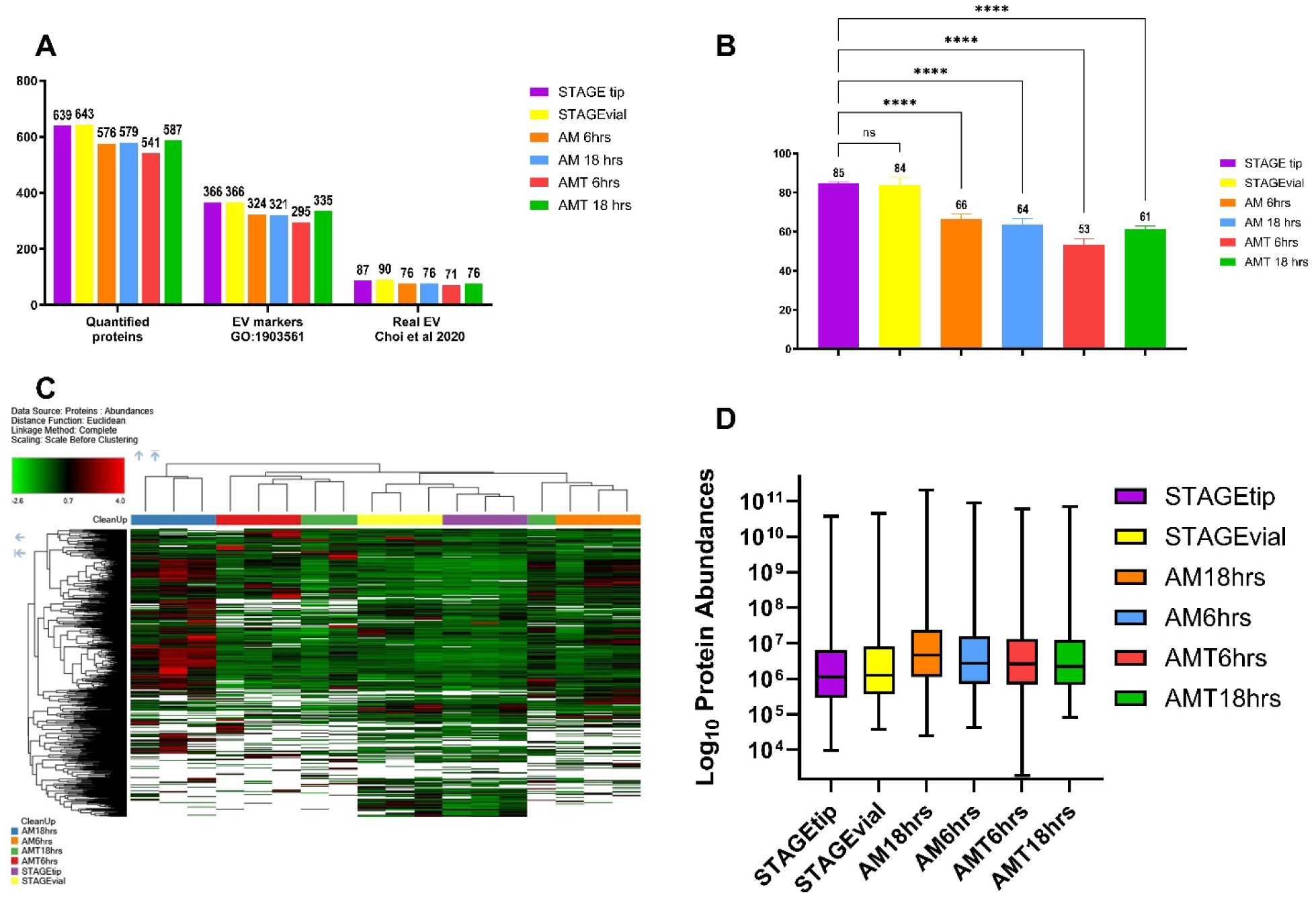
Grouped abundances plot (A), “real” EV marker number plot (B), heatmap (C) and box and whisker plot for grouped abundances for comparison of digestion buffer (AM=Ammonium Bicarbonate, AMT+ Ammonium Bicarbonate +10% TFE), digestion time (6hrs or 18hrs) without STAGEtip desalting with standard digestion (100 mM HEPES + 10% TFE, 18 hrs) and STAGEtip desalting. Additional, comparison of elution of STAGEtips directly into glass micro insert for LC-MS vials (STAGEvial). Statistical significance calculated by one-way ANOVA. P-value <0.05 (*), <0.01 (**), <0.001 (***), <0.0001 (****).

**PART 4: Confounding factors (3.9-3.10)**

Platelet contamination and sample hemolysis are both major confounding factors for EV proteomics. Section 3.9 investigated the impact of platelet contamination in single donor human to test for enrichment of PF4 as a candidate platelet marker. Additionally, we test the impact of the introduction of a filtration step into SEC-SAX protocol to remove residual platelets. Section 3.10 investigates the impact of sample hemolysis in single human donor human plasma samples and the impact of the post storage centrifugation step on residual platelet contamination. Refer to Table 2 for a summary of the preparation of plasma for Part 4.

**Table 2:** Summary of centrifugation and filtration steps in preparation of plasma for Part 4.

|  | Plasma preparation |  |  |  |  |
| --- | --- | --- | --- | --- | --- |
| Plasma | 1000 g<br>10 min | 2500 g<br>15 min | 2500 g<br>15 min | 10,000 g<br>20 min | filtration |
| **Section 3.9: platelet contamination and filtration** |  |  |  |  |  |
| Normal | √ | √ | √ | √ | - |
| Normal (filtered) | √ | √ | √ | √ | √ |
| Platelets | √ | - | - | - | - |
| Platelets (filtered) | √ | - | - | √ | √ |
| **Section 3.10: Hemolysed human plasma** |  |  |  |  |  |
| Normal | √ | √ | √ | √ | - |
| Low Heme | √ | √ | √ | √ | - |
| High Heme | √ | √ | √ | √ | - |
| Platelet contaminated<br>(normal Heme) | √ | √ | √ | √ | - |

### 3.9 The impact of platelet contamination and the effect of filtration

There is significant overlap between the platelet and EV proteomes, which makes it challenging to determine whether EV samples have been affected by platelet contamination. To address this issue, we compared a published platelet proteome (Burkhart et al., 2012) with an EV proteome (Choi et al., 2020) we selected Platelet Factor 4 (uniprot accession P02776) as a candidate marker for platelet contamination. PF4 is highly abundant in platelets and is absent from the “real EV” protein list, making it a suitable indicator. Section 3.9 compares platelet contaminated and normal plasma samples to confirm PF4 as a reliable marker of platelet contamination in SEC-SAX processed EV samples (Supplemental Data S7).

The level of platelet contamination in the plasma is impacted by the number, speed and duration of centrifugation steps used during plasma preparation. There are 4 centrifugation steps in the preparation of the plasma used for the usual the SEC-SAX protocol (Figure 18). The first is to remove the red blood cells (1000 g 10 minutes), the next two are both to remove platelets (2500 g for 15 minutes each) and then in our standard plasma protocol the samples are frozen at-80°C. Once thawed the samples are centrifuged at 10,000 g for 20 minutes to remove the larger extracellular vesicles and then samples are prepared for proteomics using the SEC-SAX protocol. To test the impact of platelet contamination, high platelet plasma samples were prepared with only a single 1000 g centrifugation step to remove red blood cells before storage. The post storage 10,000 g centrifugation step was also omitted to retain platelets. Plasma samples were prepared at the same time with the usual 4 centrifugation steps for comparison.

**Figure 18:**
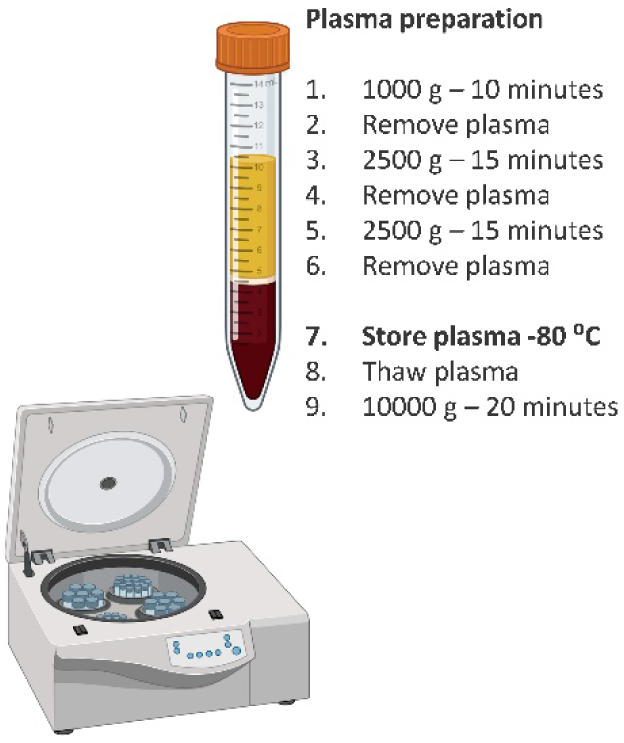
Centrifugation protocol for the preparation of high-quality plasma from blood samples for the SEX-SAX protocol.

There were 1607 proteins quantified from the platelet contaminated samples compared with 945 for normal plasma. Of these 1607 platelet contaminated proteins 45.8% (736 proteins) were categorised as EV markers by STRING analysis and 201 proteins (12.5%) as “real EV” markers. Of the 945 proteins quantified in the normal plasma, STRING analysis identified 53% (501 proteins) as EV markers and 136 (14.4%) as “real EV” markers (Figure 19A). This increased number of quantified proteins and reduced percentage of EV markers indicates that the platelet contamination impacts the quality of the proteomics data. Similar numbers of proteins were quantified from the filtered samples (Figure 19A). However, filtering the EV samples (SEC eluant prior to SAX) resulted in significant protein loses. This is seen in the box and whisker plot of protein abundances for all quantified proteins (Figure 19B). Additional experiments with filtration caused highly variable sample losses resulting in inconsistent data (Data not shown). Filtering was not included in the SEC-SAX protocol, and all further data is without filtration. There were 672 quantified proteins unique to the platelet contaminated samples (Supplemental Data S8), which included TAR DNA-binding protein 43 (gene symbol: TARDBP, Accession: Q13148), Superoxide dismutase (gene symbol: SOD1, Accession: P0044), and Alpha-synuclein (gene symbol: SNCA, Accession: P37840) which are known markers of neurodegenerative disease (Wiersema et al., 2024). Additionally, P-selectin (SELP)(CD42p) which is a marker of activated platelets was also unique to the platelet contaminated samples (Suades et al., 2022)(Figure 19C).

**Figure 19:**
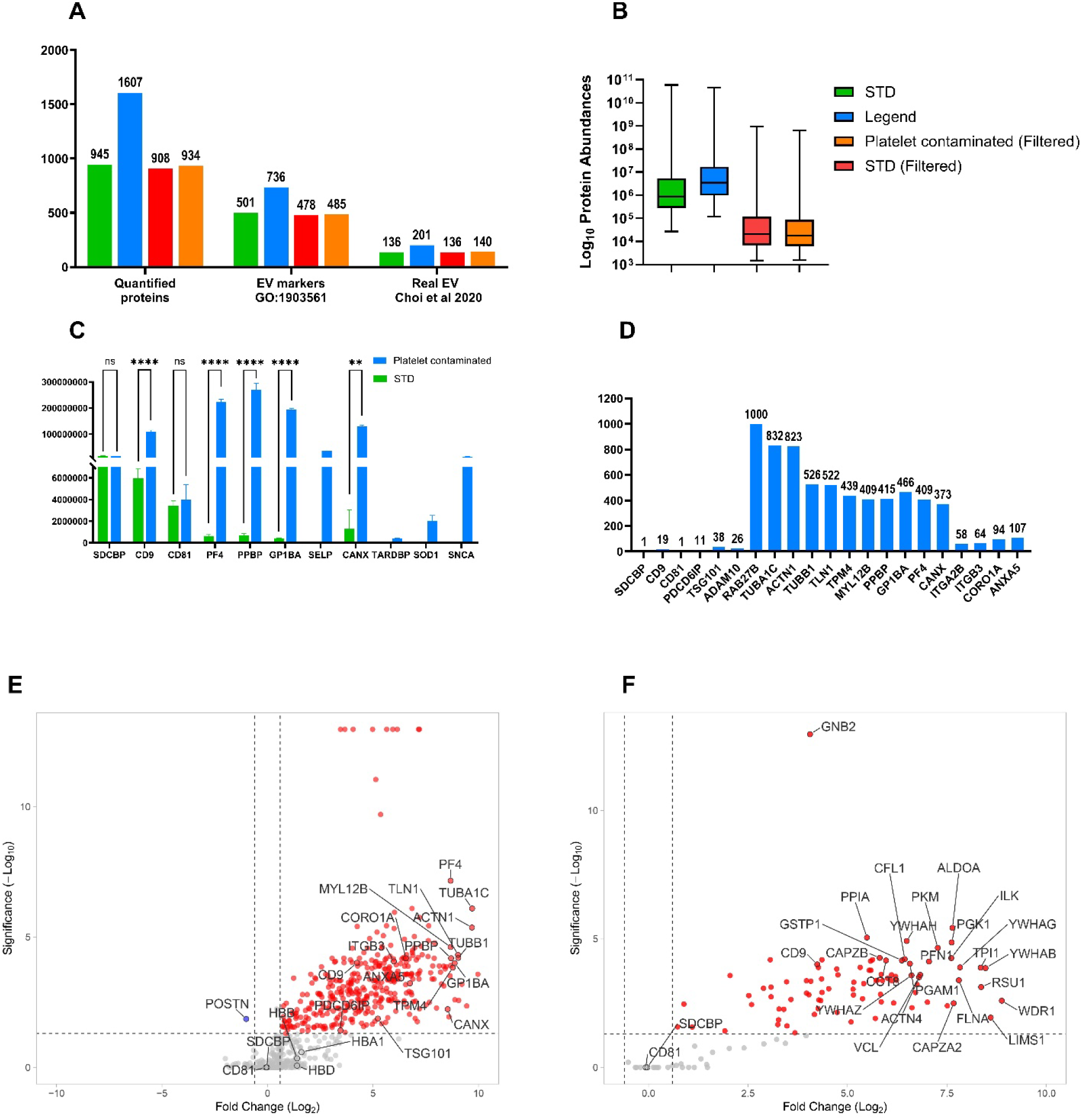
Impact of platelet contamination and filtration on the plasma EV proteome. Quantified protein in normal plasma, and platelet contaminated sample with and without filtration (Panel A). Protein abundance box plot of normal and platelet contaminated samples with and without filtration (Panel B). Protein abundance presented as Mean + SD for EV markers Syntenin, CD9 and CD81, candidate platelet markers PF4, PPBP, GP1BA and calnexin without filtration (Panel C). Fold change with platelet contamination for EV markers and candidate platelet markers (Panel D). Volcano plot comparing all quantified proteins (Panel E) from normal and plasma contaminated samples without filtration and real EV proteins (Panel F). Statistical significance calculated by Proteome Discoverer fold-change ≥1.5 and P-value <0.05 (*), <0.01 (**), <0.001 (***), <0.0001 (****) (Plot C).

There were 935 proteins quantified in both normal and platelet contaminated plasma, of which 373 proteins were significantly increased in the platelet contaminated plasma (Supplemental Data S9) (Figure 19E). This included PF4, Calnexin (CANX), CD9 but not CD81 or Syntenin-1 (Figure 19E & C). PF4 is increased 409-fold in the platelet contaminated samples, Calnexin 373-fold and CD9 19-fold (Figure 19D). Additional EV markers were included to provide a broader picture on the impact of platelet contamination on the EV proteome and TSG101, PDCD6IP (ALIX) and ADAM10 were increased 38-fold, 11-fold and 26-fold respectively (Kowal et al., 2016; Kugeratski et al., 2021). Additionally, there is enrichment of ITGA2B (CD41), ITGB3 (CD61), CORO1A and ANXA5 which are all markers of platelet derived extracellular vesicles (Suades et al., 2022). Of the 935 proteins quantified in both samples, 137 were “real EV” markers of which 89 proteins were significantly increased in the platelet contaminated samples (Figure 19F). This demonstrates that platelet contamination can significantly impact the plasma EV proteome and is a significant confounding factor.

To investigate other proteins highly enriched in the platelet contaminated samples, the data was sorted by abundance ratio of platelet contaminated samples compared with normal plasma and then top 10 platelet enriched proteins based sorted by abundance. This places PF4 8^th^ highest abundance of the highly enriched platelet proteins and places PF4 with Platelet Basic Protein (P02775, PPBP) and Platelet glycoprotein Ib alpha chain (P07359, GP1BA)(CD42b) at ∼400-fold enrichment and a high abundance of ∼2×10^8^ consistent with PF4 as a high abundance platelet marker (Figure 19C & D). Figure 19C shows the abundance of the key platelet and EV markers while Figure 19D shows the fold change of the top 10 most enriched high abundance platelet markers. PF4 has the highest rank based on estimated copy number of the enriched proteins when referring back the published platelet proteome (Rank 4^th^ P02776), Platelet Basic Protein is ranked 6^th^ and Platelet glycoprotein Ib alpha chain is ranked 177^th^ (Burkhart et al., 2012).

### 3.10 Hemolysed human plasma and residual platelet contamination

Human plasma samples with two levels of hemolysis (low (0.21 mg/ml) and high (0.77 mg/ml)) were compared with normal, non-hemolysed plasma. Additionally, platelet-contaminated plasma that had undergone the standard post-storage centrifugation at 10,000 g for 20 min was compared with normal plasma to assess how effective this centrifugation step is at reducing platelet contamination (Supplemental Data S10). Heavily hemolysed plasma had increased numbers of quantified proteins (Figure 20A), similar to the increase observed in platelet-contaminated plasma. However, post-storage centrifugation substantially reduced this effect (compare with Figure 19A). The heat map also indicates an increase in protein abundances with hemolysis and residual platelet contamination (Figure 20B). Hemolysis caused significant increases in the abundances of EV marker CD9 and platelet marker PF4, while CD81 and Syntenin-1 were not significantly affected (Figure 20C, D). In this experiment, PF4 levels in the “residual platelet contamination” samples were restored to normal following the 10,000 g post storage centrifugation (Figure 20C, D, I). The grouped abundances plot of all proteins was also more consistent with normal plasma after this centrifugation (Figure 20E). Additionally, the neurodegenerative disease-associated proteins SOD1, SNCA and TARDBP, previously elevated in platelet contaminated samples were no longer detected, indicating effective removal of residual platelets.

**Figure 20:**
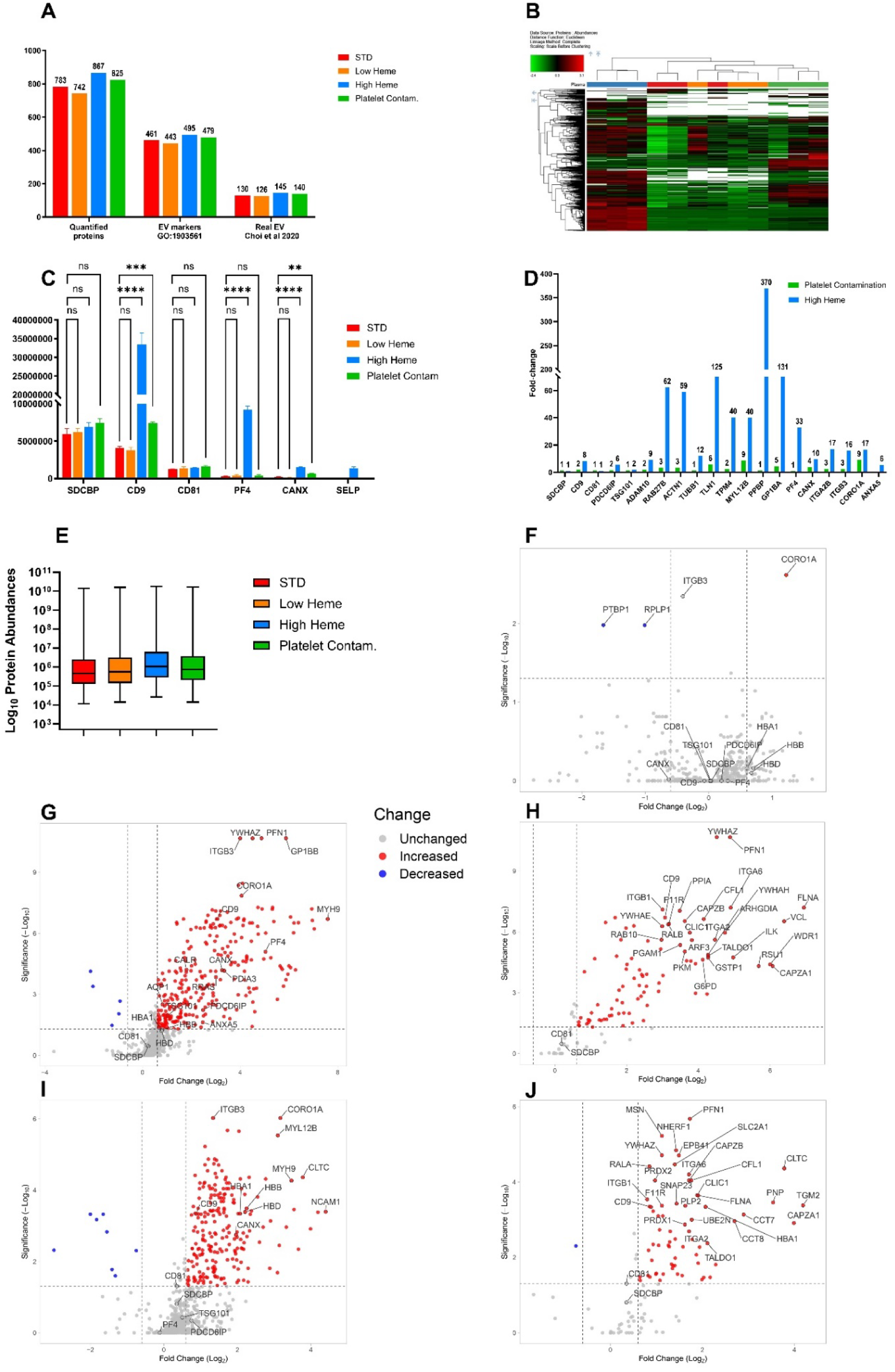
Impact of sample hemolysis and residual platelet contamination on the plasma EV proteome. Quantified proteins in normal plasma, 2 levels of sample hemolysis (low and high) and residual platelet contamination are shown based on grouped protein abundances (Panel A). Overall protein abundance distributions are presented as a box-and-whisker plot (Panel B). Protein abundance for selected EV markers and platelet markers are presented as Mean + SD (Panel C). Fold-change differences for EV and platelet markers under platelet-contaminated conditions (Panel D). Volcano plot comparing all quantified proteins (Panel F, G, I) and “Real” EV markers (H,J) from samples with low (F) and high hemolysis (G,H) and residual platelet contaminated samples (I,J). Statistical significance calculated by Proteome Discoverer fold-change ≥1.5 and P-value <0.05 (*), <0.01 (**), <0.001 (***), <0.0001 (****) (Plot C).

This highlights the importance of centrifugation in the preparation of high-quality plasma for EV purification. PF4 is a high abundance platelet marker, thus elevated PF4 reliably indicates platelet contamination. However, normal PF4 levels do not guarantee absence of platelet-derived protein contamination. PF4 elevation is therefore best considered an indicator of platelet contamination. In the “residual platelet contamination” samples, other platelet-derived markers, including Platelet glycoprotein Ib alpha chain (GP1BA) and the platelet-derived EV proteins ITGA2B, ITGB3 and CORO1A were elevated. These proteins may serve as additional indicators of plasma samples that has not been adequately prepared prior to storage.

One protein, CORO1A, was significantly increased in the low heme sample (0.21 mg/ml heme) compared to the normal plasma (0.05 mg/ml) prepared at the same time (Figure 20F). In contrast, there were 291 proteins significantly elevated (fold-change >1.5, p-value <0.05) in the high heme samples (Figure 20G), including 89 EV proteins (Figure 20H) (Supplemental Data S11).

The heavily hemolysed samples also showed evidence of platelet contamination. This was indicated by the high abundance platelet-specific markers PF4, PPBP and GP1BA, as well as the platelet-derived EV markers ITGA2B (CD41), ITGB3 (CD61), ANXA5, CORO1A (Figure 20D) and P-selectin (Figure 20C). Additional proteins that are not platelet-specific but were previously elevated in the platelet-contaminated samples (Figure 19), including CD9, CANX, PDCD6IP, TSG101, ADAM10, RAB27B, ACTN1, TUBB1, TLN1, TPM4 and MYL12B, were also increased. These further supports platelet contamination in the heavily hemolysed samples (refer to Section 3.9). TUBA1C was not detected.

The heavily hemolysed samples were also enriched for erythrocyte membrane proteins, based on comparison with the Ravinhill list of 160 proteins (Ravenhill et al., 2019). Of the 867 total proteins quantified in the heavily hemolysed samples, 65 matched proteins from the Ravinhill list. Among these, 33 were significantly enriched, including high abundance erythrocyte membrane proteins SLC4A1 (Band 3 anion transport protein), AQP1 (Aquaporin-1), BSG (Basigin) and ABCB6 (ATP-binding cassette sub-family B member 6) (Supplemental Data S13). Enrichment of erythrocyte membrane proteins was significant (Fisher exact test p=0.0038), suggesting that hemolysis leads to release of RBC-derived EVs.

In the residual platelet-contaminated samples, 223 proteins were elevated (Figure 20I), including 67 EV proteins (Figure 20J) (Supplemental Data S12).

**PART 5: Mouse plasma samples**

The SEC-SAX protocol was evaluated in mouse plasma to confirm is applicability across species. Data from eight non-hemolysed (heme ≤0.3 mg/ml), PF4-negative samples are presented as an example of SEC-SAX EV enrichment in individual mice (Supplemental Data S14). Across the eight samples, a total of 1044 proteins were quantified, and more than 60% (660 proteins) of the proteins were quantified in all samples. Mouse accession numbers were mapped to primary gene names (IDmapping) using UniProt to allow comparison with the human datasets (Supplemental Data S15). Of the quantified proteins, 554 (54%) were classified as extracellular vesicle-associated by STRING analysis (https://string-db.org/), and 156 (15%) matched the experimentally determined “real” EV marker list (Figure 21A). The GO cellular component enrichment profile from the STRING analysis (Figure 21B) was consistent with successful EV enrichment from mouse plasma. Additionally, significant enrichment of genes associated with neurological disease pathways was observed (Figure 21C).

**Figure 21:**
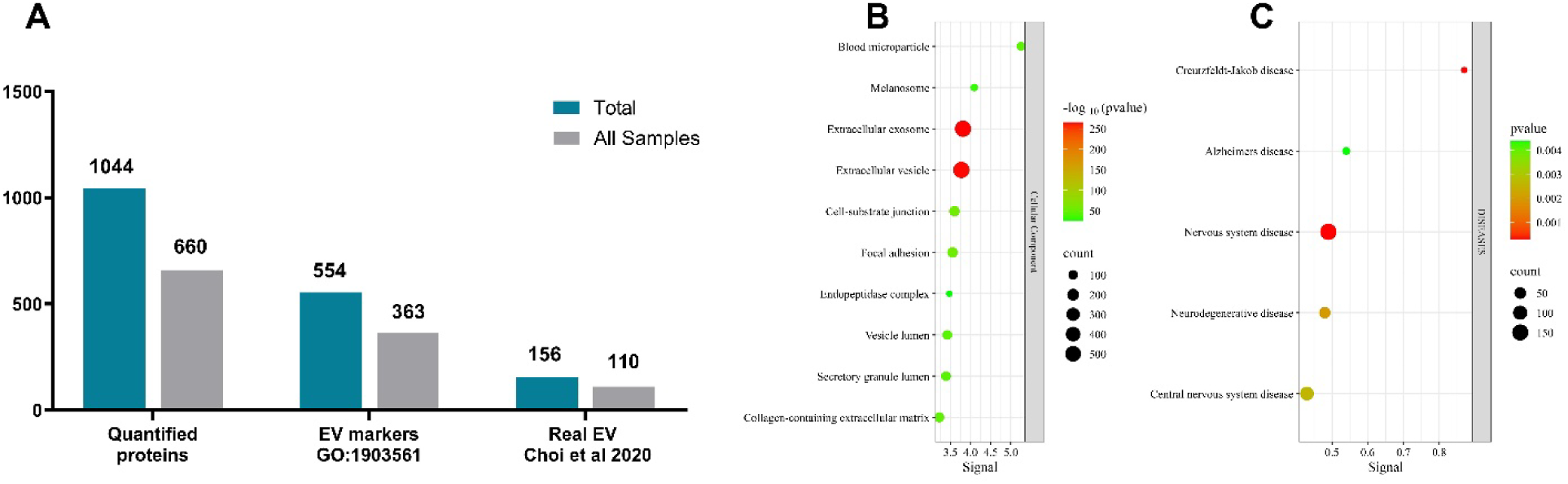
Grouped abundances for mouse plasma EV enriched by SEC SAX (n= 8 individual mice). Total quantified proteins and proteins common to all samples, analysed for number of EV proteins by STRING analysis and Choi “Real” EV markers are shown in Panel A. Gene Ontology for Cellular Component enrichment results for all 1044 quantified proteins using STRING are shown in Panel (B). Disease-associated enrichment in shown in Panel (C) (Permalink: https://version-12-0.string-db.org/cgi/network?networkId=bila16nJU6L8)(B). Visualisation of enrichment data was performed using SRplot (Tang et al., 2023).

## 4.0 Discussion

The SEC-SAX protocol was developed to specifically meet the requirements of plasma EV sample preparation from biobank samples for biomarker discovery by LC-MS-based proteomics. The aim was to develop a protocol that was applicable to both human and mouse plasma to allow for discovery studies in both preclinical mouse models and human clinical samples. The SEC-SAX protocol is highly detailed (Appendix A) and extensively tested, producing robust data from small starting volumes of plasma. The major confounding factors are specifically accounted for with quality control metrics and/or mitigation strategies. Here we describe the progressive enhancement of the SEC-SAX protocol specifically for the enrichment of small EVs from plasma for quantitative proteomics analysis. Iterative rounds of targeted protocol refinement combined with LC-MS analysis were used to improve EV purity, minimise sample loss, and reduce technical variability. The proteomics data were externally validated against both the experimentally-derived “real EV” protein list (Choi et al., 2020) and the STRING database (Szklarczyk et al., 2023). Application of the SEC-SAX protocol to 500 µl of human plasma produces data in triplicate (150 µl per replicate) quantified >600 proteins, of which >50% listed as EV-associated based on Extracellular Vesicle Gene Ontology (GO: 1903561), and 80-100 classified as “real EV” proteins including key EV markers CD9, CD81 and Syntenin. As only 150 µl of plasma is required per replicate, this workflow is suitable for mouse studies using individual animals without the need for sample pooling. Application of the SEC-SAX protocol in individual mouse plasma samples demonstrates EV enrichment and quantification of biomarker proteins associated with neurodegenerative diseases.

The final protocol can be divided into 5 main steps: **1)** Plasma preparation **2)** Size Exclusion Chromatography **3)** Strong Anion Exchange **4)** single-pot, solid phase-enhanced sample-preparation (SP3) and **5)** trypsin digestion and Stage tip desalting. During protocol development, we evaluated the contribution of each step, allowing us to remove procedures that did not improve EV enrichment or proteomics performance. Due to the modular nature of the protocol, individual steps can be modified, optimised further, or substituted with alternative workflows as needed.

### Plasma preparation

**Step 1** is the preparation of plasma by three centrifugation steps prior to storage. The sequential centrifugation steps at room temperature remove red blood cells and platelets without platelet activation. Samples stored at - 80⁰C are thawed on ice then precleared by centrifugation at 10,000 g to remove larger vesicles and residual platelets. Careful handling and consistent pipetting of samples is critical for removal of platelets, which are a major confounding factor for EV proteomics (**Section 3.9**). The level of hemolysis should be measured using a nanodrop spectrophotometer using the custom heme protocol prior to freezing where possible and hemolysed samples excluded due to the impact on the EV proteomics signature (**Section 3.10**).

### Size exclusion chromatography

**Step 2**, Size exclusion chromatography separates the EVs from the soluble plasma proteins and effectively removes >99% of albumin (**Section 3.3 Figure 7**). The precision and consistency of this step directly impacts the variability and quality of the proteomics data. Analytical pipettes are preferred over the use of a droplet based automatic fraction collector (**Section 3.3 Figure 3**). Scaling down to the 150 µl qEVsingle 35nm columns is important, as it enables processing of triplicate human samples from only 500 µl of plasma (**Section 3.5 Figure 12** samples 11-13; first use of the 150 µl qEVsingle columns and for all subsequent sections). This format also allows processing of single 150 µl samples from individual mice (**Section 3.11**).

### Strong Anion Exchange (SAX)

**Step 3**, Strong Anion Exchange (SAX) magnetic beads are used to concentrate the EVs directly from the dilute SEC eluant without prior concentration or buffer exchange (**Section 3.5**). Using Bis-Tris propane as the SEC buffer is required to avoid a decrease in the number of EV markers. Sample loss can occur with centrifugal filters by absorption of EVs to the filter membrane. Eliminating the need for centrifugal filters maintains the yield of EV proteins from small starting volumes of plasma. Additionally, anion exchange preferentially captures EVs over lipoproteins. Therefore, the incorporation of magnetic SAX beads not only reduces sample loss but also enriches for EVs over contaminating lipoproteins.

### EV lysis and Single Pot enhanced (SP3) sample preparation

**Step 4**, The EV proteins are recovered by EV lysis and protein solubilisation by boiling the SAX beads in 2% SDS (in 100 mM HEPES). Solubilisation of the EV bilayer membrane is necessary for trypsin digestion of the intravesicular proteins (Choi et al., 2020). Single Pot enhanced (SP3) sample preparation removes the SDS prior to trypsin digestion. The single-tube nature of SP3 minimises sample handling and is well known for its low sample-loss characteristics, as the magnetic beads act as carriers for proteins throughout the clean-up process (Hughes et al., 2019). The SAX beads were not able to be used for the SP3 protein aggregation capture in place of the usual mixture of hydrophobic and hydrophilic carboxylate magnetic beads (Cytiva) without the loss of CD81 from the proteomics data (**Section 3.5 samples 9-10**). It would have been preferrable to use the SAX beads for both the anion exchange capture of the EVs and the preparation of samples for trypsin digestion (Wu et al., 2025). This would have reduced the number of sample transfers and eliminated the need for the carboxylate beads. However, we considered the loss of CD81, a canonical EV marker, unacceptable.

### Trypsin Digestion and desalting

**Step 5**, The use of a non-volatile HEPES digestion buffer necessitates a post digest desalting step. The post digest desalting step is often omitted when SP3 is used to clean up samples prior to digestion as long as a volatile digest buffer that can be removed by evaporation is used (Hughes et al., 2019). We observed that including the STAGEtip solid phase extraction interestingly increased EV proteome depth of the low-abundance EV proteins, even though it resulted in greater overall sample loss compared with digestion in ammonium bicarbonate followed by evaporation (**Section 3.8**). Additionally, the STAGE tips also act as a secondary method to remove any residual magnetic beads from the SP3 step. These beads can damage or block the nano LC-MS columns and microfluidics (Kanao & Ishihama, 2025; Rappsilber et al., 2003). Although the use of individual STAGE tips increases manual handling, their contribution to deeper EV proteome coverage and instrument protection made them essential, and they were therefore retained in the final protocol.

Direct comparison between proteomics studies is challenging due to differences in mass spectrometer sensitivity, data acquisition and data processing methods in addition to the sample preparation (Guo et al., 2025; Singh et al., 2025; Vallejo et al., 2023). For this reason, our paper focuses on relative EV purity and EV specificity, assessed using internal controls and external reference datasets. We assess our proteomics data quality in the context of the three main confounding factors for plasma-derived extracellular vesicles, namely lipoproteins, sample hemolysis and platelet contamination (Lucien et al., 2023; Welsh et al., 2024). EV purity ratios were devised as an indication of enrichment of EVs over plasma contaminants (**Section 3.3, 3.6-3.7**). We selected albumin and ApoB as representative markers of soluble plasma proteins and lipoproteins, respectively, as they were the most abundant members of these contaminant classes. The same principle can be applied to other proteins such as IgG, as an indication of immunoglobulin contamination, or PF4, as an indication of platelet contamination. We chose an objective measure of sample hemolysis, as qualitative assessment of hemolysis by eye is unreliable. We then tested the impact of hemolysis on the proteomics data (**Section 3.10**). Interestingly, hemolysed samples show evidence of platelet contamination and potentially both erythrocyte and platelet derived EVs. As evidenced by platelet EV markers and red blood cell membrane proteins respectively. This suggests that hemolysis may not simply be the rupture of red blood cells, but potentially the activation and release of blood cell derived EVs due to shear stress (S. R. Ma et al., 2023; Suades et al., 2022). Platelet contamination without sample hemolysis is more difficult to discern, which is further complicated by the extensive overlap between the platelet and EV proteomes (**Section 3.9**). We selected Platelet Factor 4 (PF4) as a candidate platelet marker due to its high abundance in platelets (Burkhart et al., 2012) and absence from the real EV list (Choi et al., 2020). We then confirmed its relative enrichment in platelet contaminated samples (**Section 3.9**). PF4 was enriched 409-fold in platelet-contaminated samples. However, the biological function of PF4 and its known release from activated platelets is potentially more important as elevated PF4 in a sample cohort suggests that the proteomics signature is potentially due to platelet contamination and/or of platelet origin. PF4 has previously been identified as a differentially enriched EV protein from plasma samples and suggested as a potential disease biomarker. Our findings therefore raise questions about the interpretation of those datasets (Aparicio et al., 2024; W. Ma et al., 2020) and additional studies reporting PF4-associated EV signatures (Su et al., 2024; H. Yin et al., 2024a). Platelet contamination has significant implications for biomarker discovery studies that rely on circulating EVs as surrogates for tissue sampling. Contaminating platelet proteins can lead to false-positive identification of biomarkers of disease, as evidenced by TAR DNA-binding protein 43 (TARDBP; Q13148), superoxide dismutase 1 (SOD1; P00441), and alpha-synuclein (SNCA; P37840), which are established neurodegenerative disease markers (Wiersema et al., 2024) that are also present in normal platelets (Section 3.9). For this reason, we tested the addition of a 0.2 µm filtration step between step 2 (SEC) and step 3 (SAX) on platelet contaminated samples. However, filtration caused significant sample loss and increased variability (**Section 3.9**). These effects undermine reproducibility, reduce signal-to-noise and can obscure the EV proteomic signature; therefore, we did not include filtration in the SEC-SAX protocol. In contrast, the standard post-storage 10,000 g centrifugation step reduced, but did not eliminate, residual platelet contamination. Additional platelet-associated proteins may be valuable as quality-control markers to assess plasma preparation, including Platelet Basic Protein (P02775; PPBP) and Platelet glycoprotein Ib alpha chain (P07359; GP1BA) (Burkhart et al., 2012) and platelet EV markers P-SELP, ITGA2B, ITGB3, CORO1A and ANXA5 (Suades et al., 2022). Together, these objective Quality Control (QC) measures and quantitative effects on the EV proteome help address the under-quantified impact of key confounding factors in LC-MS-based EV studies.

Data-dependent acquisition (DDA), which prioritises the most abundant peaks, was used in the development of the SEC-SAX protocol to highlight dominant proteins. While this approach reliably quantified >600 proteins per sample from small plasma inputs, advanced acquisition strategies such as data-independent acquisition (DIA) and FAIMS could further increase EV proteome depth and realise the full potential of this workflow. These complementary strategies can be layered onto SEC-SAX without altering the sample preparation or protocol. Post-acquisition data normalisation and cleaning should also be explored, as the choice of normalisation and analysis strategy will be critical to counteract inherent EV sample heterogeneity. In conclusion, we present a modular SEC-SAX protocol for enriching EVs from small plasma volumes, applicable for human and mouse studies. Incorporating objective hemolysis screening and PF4-based platelet QC measures ensures high-purity EV proteomes for reliable biomarker discovery.

## Supporting information

Graphical Abstract

## Acknowledgements

This work was supported by funding provided by Institute for Physical Activity and Nutrition, School of Exercise and Nutrition Sciences and FightMND under Grant IP-158.

## Data availability

Proteome Discoverer results are provided as Excel exports in SupplementalData.xlsx, and the figure source data (including STRING permalinks) are provided in SourceData.xlsx; raw mass spectrometry data will be deposited in the PRIDE repository upon acceptance of the manuscript.

## Appendix A: Detailed SEC-SAX Protocol

### Materials and Equipment Setup

**1. SEC Column Equilibration:** Remove the Size Exclusion Chromatography (SEC) columns from 4°C storage and allow them to equilibrate at room temperature for at least 1 hour prior to use.

- *Note:* SEC columns are stored at 4°C to prolong resin stability and prevent cracking of the stationary phase.
- *Note:* A batch of 24 can be handled efficiently by a single person
**2. Buffer Preparation:** Prepare the **Bis-Tris Propane (BTP) SEC Buffer** and **SAX Bead Buffer** fresh for each experiment to prevent microbial growth and ensure optimal pH stability. Buffers must be sterile filtered to remove all particulate matter. Filter the buffers into sterile 50 ml Falcon tubes using a separate 0.2 µm PES syringe filter and a 10 ml syringe for each buffer to prevent cross-contamination.
**3. Refer to Table 3 (pages 8-9) for additional reagents**

### Step-by-step method details

#### Plasma Preparation: pre-clearing from frozen aliquots

##### Timing: ∼1.5 hours on plasma aliquot volume and number of samples

Note: While plasma is thawing, equilibrate SEC columns and SAX beads as detailed in the subsequent sections.

**1. Thawing:** Thaw plasma aliquots on ice. Warm the tubes by hand and mix by inversion until cryoprecipitates are fully dissolved.
a. *Note:* Larger aliquots (∼1 ml) may take up to an hour to thaw. It only takes a few minutes to dissolve the cryoprecipitates, which appear as cloudy material in the yellow plasma. The plasma becomes clear once these precipitates have fully dissolved.
**2. Centrifugation:** Centrifuge the thawed plasma at **10 000 g, 20 min, 4°C**. Ensure the tube hinge faces upward to orient the pellet.
**3. Supernatant Recovery:** Using a P200 pipette, carefully transfer the supernatant (pre-cleared plasma) to a fresh 1.5 ml Eppendorf tube. Avoid contact with the pellet located on the hinge side; while the pellet contains residual platelets and larger vesicles, it is usually not visible.
`o` *Small Volume Handling:* For small-volume samples (e.g., mouse plasma), leave an absolute minimum of **10 μl** residual volume in the original tube. Aim for consistency in this residual amount across all samples.
`o` Keep the pre-cleared plasma on ice until further processing.

#### Preparation of SAX Beads

##### Timing: 30-45 minutes

1. **Resuspension:** Swirl the **MagReSyn® Strong Anion Exchange (SAX)** bead stock using a circular motion to mix thoroughly and ensure a homogeneous suspension.
2. **Aliquoting:** Aliquot **20 μl** of SAX beads into individual 1.5 ml Eppendorf tubes. Mix the bead suspension between each aliquot to maintain uniform suspension.

`o` *Note:* Swirling the bottle 3-5 times between each aliquot is usually sufficient, provided enough force is used.

**CRITICAL:** Consistent aliquoting of the SAX beads is essential for reproducible results, as the bead volume directly impacts EV recovery. The beads settle quickly; vigorous swirling of the stock bottle is necessary between dispensing aliquots.

`o` *Bulk Option:* Alternatively, beads can be washed as a pool. 2 ml Low-bind Eppendorf tubes are recommended for bulk washing to prevent bead loss on the tube surface. SAX beads in 2 ml tubes can be mixed efficiently by inversion. Usually, 3 inversions are sufficient for resuspension between each aliquot. A flick gesture can be used to get the beads off the lid of the tube during aliquoting.
**3. Equilibration:** Pulse spin beads (500 rpm, 30 seconds) to remove any liquid from the tube lid and place aliquoted beads on a magnetic rack for 3 minutes. Check that the bead volume is consistent between tubes.
`o` Remove storage buffer with a P200 pipette and add **600 µl of SAX Bead Buffer (BB: 50 mM Bis-Tris Propane, pH 6.5, 150 mM NaCl)**. A repeater pipette with a 10 ml Combitip can be used for efficiency.
`o` Equilibrate beads on a rotary wheel for 5 min at 4°C.

Recover beads on the magnetic rack as before (spin beads 500 rpm, 30 seconds and then 3 minutes on magnetic rack); remove the wash buffer slowly with a P1000 pipette to prevent bead loss, and discard buffer.

**4. Hold step:** Keep tube lids closed between washes to prevent bead drying. Beads should be washed twice and kept in Bead Buffer until immediately before EV elution.
`o` *Note:* Once equilibrated, tubes can remain on the magnetic rack ready for buffer removal immediately prior to SEC elution.

#### SEC Column Equilibration (Bulk)

**Timing:** ∼1 hour (Typical flow rate is 10 min per 3 ml column volume).

**1. Setup:** Arrange all columns in racks and equilibrate as a batch. Ensure columns have reached room temperature prior to BTP SEC buffer equilibration.
**2. Storage Buffer Removal:** Remove caps in order: first the top blue cap, then the bottom cap. Allow storage buffer to drain completely until the upper frit is just exposed to air.
**3. Flushing:** Fill columns with **3 ml of filtered BTP-SEC buffer**. (The head space holds 3 ml; a repeater pipette set to 1.5 ml x 2 is efficient here). Flush columns with a total of **9 ml (3 column volumes)** of BTP-SEC buffer (pH 6.3). Keeping the columns topped up decreases flush time.
**4. Stopping:** Stop columns with ∼1 cm of buffer remaining above the frit by capping the bottom. This allows slower columns to catch up so all samples can be loaded in a similar timeframe. Capping should be done in reverse order (bottom cap first).

#### EV Size Exclusion Chromatography

**1. Batching:** Process columns in batches of 6 to minimize variation due to sample diffusion. Alternate samples and controls to prevent order effects.
**2. Loading Preparation:** Remove the bottom cap and run out the remaining 1 cm of buffer immediately prior to loading. To avoid plasma dilution, the column should stop dripping and the frit should be just exposed, but do not let columns sit for more than a few minutes without buffer.
**3. Sample Application:** Apply **150 μl** of pre-cleared plasma directly to the frit using a P200. Ensure a clean layer of plasma without drips or splashes on the column walls.
**4. Void Volume:** Wait until the sample has completely entered the resin bed (indicated by a dry frit and stopping of dripping). Then add **550 μl of BTP-SEC buffer** directly to the frit. Apply buffer in a smooth, slow flow to avoid disturbing the plasma layer.

- *Note:* The total void volume of **700 μl** (150 µl plasma + 550 µl buffer) is discarded.
**5. Preparation for Elution:** While the 550 µl void is running, remove the buffer from the equilibrated SAX beads. Use a clean Kimwipe to remove any residual drops of void from the SEC column tip once the drips stop.
**6. EV Elution:** Elute EVs directly into the tubes containing the equilibrated SAX beads in a single fraction of **680 μl BTP-SEC buffer**.
**7. Overnight Incubation:** Place eluted EVs on ice and add **6.8 μl protease inhibitors**. Once all batches (up to 24 samples) are processed, seal lids with parafilm and incubate on a suspension mixer **overnight at 4°C**.
**8. Bead buffer storage:** store buffer overnight at 4⁰C ready for bead washing the next day

**Table 1:**
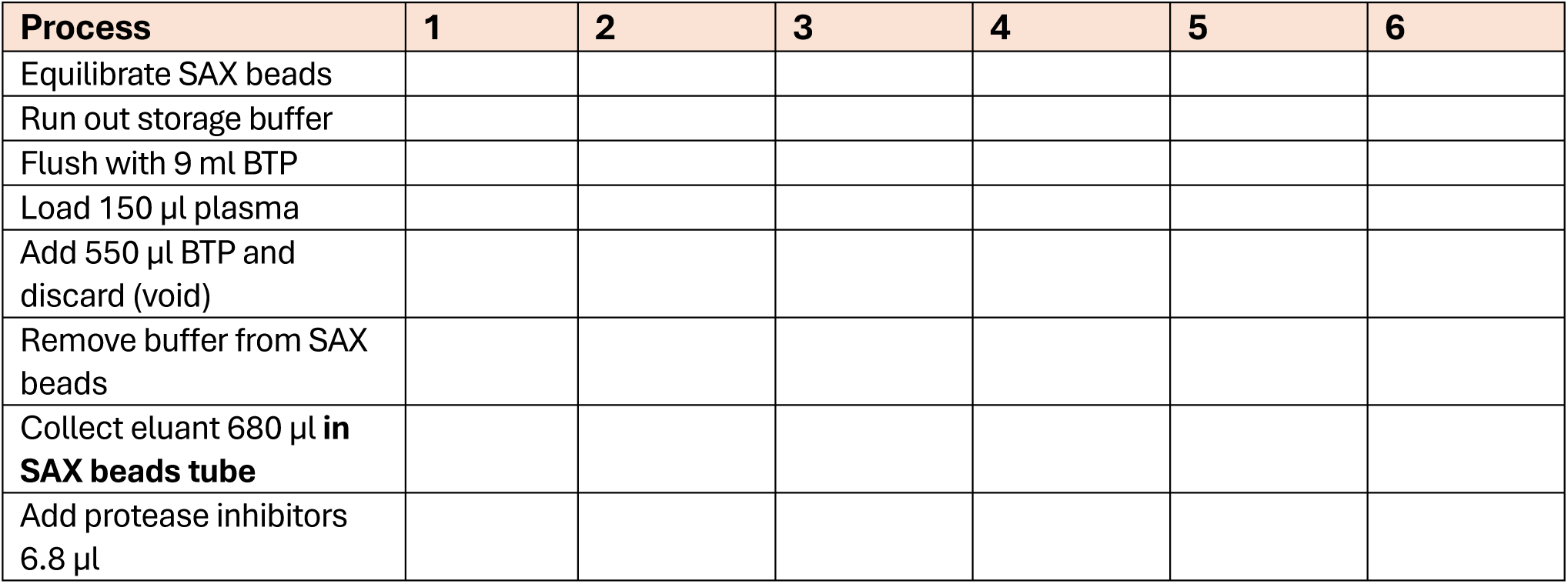
SEC checklist.

#### Bead Washing and Lysis (Day 2)

1. **Washing:** Capture beads magnetically (spin beads 500 rpm, 30 seconds and then 3 minutes on magnetic rack). Wash **3 times** with **900 μl ice-cold wash buffer**. Incubate for 5 min at 4°C on a suspension mixer for each wash. Remove wash buffer slowly with a P1000 pipette.
2. **Lysis:** During the final wash, prepare Lysis Buffer (Table 1). Add **200 μl EV Lysis Buffer (2% SDS, 100 mM HEPES, pH 7.5)** to each tube.

- *Note:* ResynBio SAX beads tend to stick to the inside of the tube especially with human plasma. Mouse plasma which contains less ApoB is much less sticky. To recover proteins, vortex extensively then rotate 1.5 ml tubes (nested in 50 ml Falcons) on a tube roller for two 5-min intervals at 4⁰C. Vortex extensively before and after each interval on the tube roller.
3. **Heating:** For complete lysis heat samples at **95°C for 15 min** using a Thermomixer without mixing. Allow to cool to room temperature by resetting the Thermomixer to 22⁰C.

- **Note:** samples should not be mixed during heating or removed from the Thermomixer above 50⁰C as the tube lids can pop open due to vapour pressure resulting in loss of sample volume.
4. **Recover EV lysate:** Capture beads magnetically (spin beads 6000 rpm, 30 seconds and then 3 minutes on magnetic rack) and transfer the EV lysate (supernatant) to new 1.5 ml tubes. Discard the tubes with spent SAX beads.

### Mass Spectrometry Sample Prep: Reduction, Alkylation, and SP3

1. Reduction and Alkylation:

`o` Add **3.6 µl 0.5M TCEP**, vortex briefly then centrifuge 6000 rpm, 30 seconds to remove liquid from the tube lid.
`o` Cover tube with foil to protect the TCEP from light
`o` Prepare the 0.5M IAA
`o` Add **16 µl of freshly prepared IAA** (0.5 M) vortex briefly then centrifuge 6000 rpm, 30 seconds to remove liquid from the tube lid.
`o` Heat at **95°C for 15 min**, then cool to 22°C using a Thermomixer same as for EV lysis.
2. SP3 Cleanup:

`o` Add **20 µl diluted and washed SP3 beads** and briefly vortex and centrifuge 500 rpm, 30 seconds.
`o` Add **240 µl 100% Ethanol** (1:1 ratio). Shake at 1600 rpm, 8 min using Thermomixer.
3. Wash:

`o` Capture beads magnetically (spin beads 500 rpm, 30 seconds and then 3 minutes on magnetic rack).
`o` Remove and discard supernatant using P200
`o` Remove tubes from magnetic rack and add **900 µl 80% ethanol**. A 10 ml Combitip can be used for efficiency.
`o` Vortex very briefly (just long enough to unstick beads from wall of tube) and mix using a Thermomixer 3 min at room temperature 1600 rpm.
`o` Capture beads magnetically (spin beads 500 rpm, 30 seconds and then 3 minutes on magnetic rack).
`o` Remove wash buffer slowly with a P1000 pipette and discard.
`o` Wash a total of **3 times with 900 µl each of 80% Ethanol**.
`o` Remove any residual ethanol wash with a P200 pipette after the final wash.

#### **CRITICAL:** SP3 beads dry rapidly and become sticky. Keep lids closed and work quickly

**4. Digestion:** Add **50 µl digestion buffer** (100 mM HEPES + 10% TFE) and resuspend each sample with P100 or P200 pipette

`o` The beads will be stuck to the side of the tube.
`o` Work quickly to resuspend the beads in the digestion buffer by washing the beads down to the bottom of the tube. The beads will stick to pipette tip if overworked.
5. **Add 10 µl diluted Trypsin** (0.01 µg/µl).

`o` *Note:* Trypsin is added separately to minimise wastage and diluted to allow for accurate pipetting
**6.** Incubate at **37°C for 18 hours** at 1600 rpm in a Thermomixer.

#### STAGE tip Solid Phase Extraction and Resuspension (Day 3)

**1. STAGEtip Setup:** Prepare SDB-RPS STAGE tips with 2 layers of membrane in a P200 pipette tip
1. These can be prepared in advance
**2.** Capture beads using magnetic rack (centrifuge max speed, 30 seconds then magnetic rack > 3 mins).
**3. Equilibration:** For each STAGEtip, add 50 µl of each solvent in order and centrifuge for the indicated time at 500 g after each addition.

`o` Acetonitrile (3 mins)
`o` 30% Methanol, 0.2% TFA (6 mins)
`o` 0.2% TFA (3 mins)
`o` *Note:* Check each STAGE tip after each centrifugation step to confirm that no solution remains in the tip. Centrifuge for an additional minute if necessary.
**4. Loading:** Add 150 µl 1% TFA to tips, layer on peptides without beads, and centrifuge at 1000 x g for 8 mins.
**5. Washing:** Wash with 100 µl 0.2% TFA (2x) and 40 µl 90% Isopropanol,1% TFA (1x) and centrifuge **1000 g, 5 mins** after each addition.
**6. Elution:** Insert a glass insert into a new 1.5 mL Eppendorf tube (ensure spring is removed). Add **60 µl of 80% acetonitrile containing 5% ammonium hydroxide**.

#### **CRITICAL:** Ammonium hydroxide is extremely volatile; handle in a fume hood. Combitips or positive displacement pipettes are recommended

`o` Centrifuge at 1000 × g for 3 minutes.
**7. Drying:** Vacuum dry at 45°C (45-60 minutes)
`o` Pause point: dried peptides can be stored indefinitely at-80⁰C
**8.** Resuspension: **Add** 10 µl of Loading Buffer A.
`o` Remove the glass insert containing dried peptide using tweezers.
`o` Add 10 µl of Loading Buffer A using a P10 pipette

**Table 2:** Solid Phase Extraction Checklist.

| Solvent | Amount | Centrifugation | Check List |
| --- | --- | --- | --- |
| **Equilibration** |  |  |  |
| acetonitrile | 50 µl | 500 g, 3 mins |  |
| 30% methanol + 0.2% TFA | 50 µl | 500 g, 6 mins |  |
| 0.2% TFA | 50 µl | 500 g, 3 mins |  |
| **Loading** |  |  |  |
| 1% TFA | 150 µl | - |  |
| Peptide | All | 1000 g, 8 mins |  |
| **Wash** |  |  |  |
| 1. 0.2% TFA | 100 µl | 1000 g, 5 mins |  |
| 2. 0.2% TFA | 100 µl | 1000 g, 5 mins |  |
| 3. 90% isopropanol + 1% TFA | 40 µl | 1000 g, 5 mins |  |
| **Elution** |  |  |  |
| 80% acetonitrile/5% ammonium hydroxide | 60 µl | 1000 g, 3 mins |  |

#### CRITICAL: Do not resuspend the peptide; drop the Loading Buffer A into the insert carefully without touching the pipette to the peptide

`o` Twist the spring back onto the bottom of the glass insert.
`o` Centrifuge at 1000 rpm, 30 seconds.
`o` Place in a sonicating water bath for 20 minutes using tube float.
`o` Centrifuge again at 1000 rpm, 30 seconds to remove air bubbles.
`o` Transfer the glass insert with resuspended peptides to a glass vial and twist on a new cap.

**Table 3:** Reagent list.

| Reagent | Process | Section | Composition | Storage |
| --- | --- | --- | --- | --- |
| SEC buffer | Mobile phase for SEC | Size Exclusion Chromatography | 50 mM Bis-Tris Propane, 75 mM NaCl, pH 6.3 | Make up fresh on day and 0.2 µm filter |
| Bead buffer | Equilibration and Washing SAX beads | SAX beads | 50 mM Bis-Tris Propane, 150 mM NaCl, pH 6.5 | Make up fresh on day and 0.2 µm filter |
| SAX beads | SAX | SAX binding SEC eluant | 20 µl equilibrated I twice in 600 µl bead buffer | Make up fresh on day |
| Lysis buffer | Elution of SAX beads | SAX beads | 2% SDS, 100 mM HEPES, pH 7.5 | Can be made up in advance, keep indefinitely at room temperature |
| TCEP | Reduction of proteins | MS prep | 0.5 M TCEP | Store aliquoted at -20°C |
| IAA | Alkylation of proteins | MS prep | 0.5 M IAA | Make up immediately before use |
| SP3 beads | Clean up of proteins for digestion | MS prep | 5 mg (100 µl) of each bead type diluted and washed in 1 ml LC-MS H<sub>2</sub>O | Can be washed in advance, stored at 4°C |
| Digestion buffer | Trypsin digestion | MS prep | 100 mM HEPES pH 7.5, 10% TFE | Make up immediately before use |
| Trypsin | Trypsin digestion | MS prep | Dilute 0.1 µg/µl stock 10-fold in | Diluted to 0.1 µg/µl in |
|  |  |  | digestion buffer immediately before digestion and add 10 µl to each digest | 50 mM LC-MS grade acetic acid, store aliquoted at -20°C |
| 30% methanol + 0.2% TFA | STAGE tip Equilibration | SPE desalting | 30% methanol + 0.2% TFA | Make up in advance, store at room temperature |
| 0.2% TFA | STAGE tip Equilibration and washing | SPE desalting | 0.2% TFA | Make up in advance, store at room temperature |
| 1% TFA | STAGE tip loading | SPE desalting | 1% TFA | Make up in advance, store at room temperature |
| 90% isopropanol + 1% TFA | STAGE tip washing | SPE desalting | 90% isopropanol + 1% TFA | Make up in advance, store at room temperature |
| 80% acetonitrile, 5 % Ammonium Hydroxide | STAGE tip elution | SPE desalting | 80% acetonitrile, 5 % Ammonium Hydroxide | Make up immediately before use |
| MS Loading buffer A | Peptide Resuspension | MS prep | 2% Acetonitrile, 0.1% TFA | Make up in advance, store at room temperature |

## References

Aparicio, P., Navarrete-Villanueva, D., Gomez-Cabello, A., Lopez-Royo, T., Santamaria, E., Fernandez-Irigoyen, J., Ausin, K., Arruebo, M., Sebastian, V., Vicente-Rodriguez, G., Osta, R., & Manzano, R. (2024). Proteomic profiling of human plasma extracellular vesicles identifies PF4 and C1R as novel biomarker in sarcopenia. J Cachexia Sarcopenia Muscle, 15(5), 1883–1897. 10.1002/jcsm.13539

Burkhart, J. M., Vaudel, M., Gambaryan, S., Radau, S., Walter, U., Martens, L., Geiger, J., Sickmann, A., & Zahedi, R. P. (2012). The first comprehensive and quantitative analysis of human platelet protein composition allows the comparative analysis of structural and functional pathways. Blood, 120(15), e73–82. 10.1182/blood-2012-04-416594

Choi, D., Go, G., Kim, D. K., Lee, J., Park, S. M., Di Vizio, D., & Gho, Y. S. (2020). Quantitative proteomic analysis of trypsin-treated extracellular vesicles to identify the real-vesicular proteins. J Extracell Vesicles, 9(1), 1757209. 10.1080/20013078.2020.1757209

Coumans, F. A. W., Brisson, A. R., Buzas, E. I., Dignat-George, F., Drees, E. E. E., El-Andaloussi, S., Emanueli, C., Gasecka, A., Hendrix, A., Hill, A. F., Lacroix, R., Lee, Y., van Leeuwen, T. G., Mackman, N., Mager, I., Nolan, J. P., van der Pol, E., Pegtel, D. M., Sahoo, S.,…Nieuwland, R. (2017). Methodological Guidelines to Study Extracellular Vesicles. Circ Res, 120(10), 1632–1648. 10.1161/CIRCRESAHA.117.309417

Dong, L., Zieren, R. C., Horie, K., Kim, C. J., Mallick, E., Jing, Y., Feng, M., Kuczler, M. D., Green, J., Amend, S. R., Witwer, K. W., de Reijke, T. M., Cho, Y. K., Pienta, K. J., & Xue, W. (2020). Comprehensive evaluation of methods for small extracellular vesicles separation from human plasma, urine and cell culture medium. J Extracell Vesicles, 10(2), e12044. 10.1002/jev2.12044

Du, S., Guan, Y., Xie, A., Yan, Z., Gao, S., Li, W., Rao, L., Chen, X., & Chen, T. (2023). Extracellular vesicles: a rising star for therapeutics and drug delivery. J Nanobiotechnology, 21(1), 231. 10.1186/s12951-023-01973-5

Garza, A. P., Wider-Eberspacher, E., Morton, L., van Ham, M., Pallinger, E., Buzas, E. I., Jansch, L., & Dunay, I. R. (2024). Proteomic analysis of plasma-derived extracellular vesicles: pre-and postprandial comparisons. Sci Rep, 14(1), 23032. 10.1038/s41598-024-74228-4

Goedhart, J., & Luijsterburg, M. S. (2020). VolcaNoseR is a web app for creating, exploring, labeling and sharing volcano plots. Sci Rep, 10(1), 20560. 10.1038/s41598-020-76603-3

Greening, D. W., Rai, A., & Simpson, R. J. (2024). Extracellular vesicles-An omics view. Proteomics, 24(11), e2400128. 10.1002/pmic.202400128

Guo, T., Steen, J. A., & Mann, M. (2025). Mass-spectrometry-based proteomics: from single cells to clinical applications. Nature, 638(8052), 901–911. 10.1038/s41586-025-08584-0

Heinzelman, P. (2018). Magnetic Particle-Based Immunoprecipitation of Nanoscale Extracellular Vesicles from Biofluids. In T. Patel (Ed.), Extracellular RNA: Methods and Protocols (pp. 85–107). Springer New York. 10.1007/978-1-4939-7652-2_8

Holcar, M., Kanduser, M., & Lenassi, M. (2021). Blood Nanoparticles - Influence on Extracellular Vesicle Isolation and Characterization. Front Pharmacol, 12, 773844. 10.3389/fphar.2021.773844

Hughes, C. S., Moggridge, S., Muller, T., Sorensen, P. H., Morin, G. B., & Krijgsveld, J. (2019). Single-pot, solid-phase-enhanced sample preparation for proteomics experiments. Nat Protoc, 14(1), 68–85. 10.1038/s41596-018-0082-x

Jeppesen, D. K., Zhang, Q., & Coffey, R. J. (2024). Extracellular vesicles and nanoparticles at a glance. J Cell Sci, 137(23). 10.1242/jcs.260201

Ji, X., Huang, S., Zhang, J., Bruce, T. F., Tan, Z., Wang, D., Zhu, J., Marcus, R. K., & Lubman, D. M. (2021). A novel method of high-purity extracellular vesicle enrichment from microliter-scale human serum for proteomic analysis. Electrophoresis, 42(3), 245–256. 10.1002/elps.202000223

Kanao, E., & Ishihama, Y. (2025). StageTip: a little giant unveiling the potential of mass spectrometry-based proteomics. Anal Sci, 41(5), 667–675. 10.1007/s44211-025-00749-1

Khanabdali, R., Mandrekar, M., Grygiel, R., Vo, P.-A., Palma, C., Nikseresht, S., Barton, S., Shojaee, M., Bhuiyan, S., Asari, K., Belzer, S., Ansari, K., Coward, J. I., Perrin, L., Hooper, J., Guanzon, D., Lai, A., Salomon, C., Kershner, K.,…Rice, G. (2024). High-throughput surface epitope immunoaffinity isolation of extracellular vesicles and downstream analysis. Biology Methods and Protocols, 9(1). 10.1093/biomethods/bpae032

Korff, K., Muller-Reif, J. B., Fichtl, D., Albrecht, V., Schebesta, A. S., Itang, E. C. M., Winter, S. V., Holdt, L. M., Teupser, D., Mann, M., & Geyer, P. E. (2025). Pre-analytical drivers of bias in bead-enriched plasma proteomics. EMBO Mol Med, 17(11), 3174–3196. 10.1038/s44321-025-00309-0

Kowal, J., Arras, G., Colombo, M., Jouve, M., Morath, J. P., Primdal-Bengtson, B., Dingli, F., Loew, D., Tkach, M., & Thery, C. (2016). Proteomic comparison defines novel markers to characterize heterogeneous populations of extracellular vesicle subtypes. Proc Natl Acad Sci U S A, 113(8), E968–977. 10.1073/pnas.1521230113

Ku, Z., Xie, X., Hinton, P. R., Liu, X., Ye, X., Muruato, A. E., Ng, D. C., Biswas, S., Zou, J., Liu, Y., Pandya, D., Menachery, V. D., Rahman, S., Cao, Y. A., Deng, H., Xiong, W., Carlin, K. B., Liu, J., Su, H.,…An, Z. (2021). Nasal delivery of an IgM offers broad protection from SARS-CoV-2 variants. Nature, 595(7869), 718–723. 10.1038/s41586-021-03673-2

Kugeratski, F. G., Hodge, K., Lilla, S., McAndrews, K. M., Zhou, X., Hwang, R. F., Zanivan, S., & Kalluri, R. (2021). Quantitative proteomics identifies the core proteome of exosomes with syntenin-1 as the highest abundant protein and a putative universal biomarker. Nat Cell Biol, 23(6), 631–641. 10.1038/s41556-021-00693-y

Lattmann, E., Rass, L., Tognetti, M., Gomez, J. M. M., Lapaire, V., Bruderer, R., Reiter, L., Feng, Y., Steinmetz, L. M., & Levesque, M. P. (2024). Size-exclusion chromatography combined with DIA-MS enables deep proteome profiling of extracellular vesicles from melanoma plasma and serum. Cell Mol Life Sci, 81(1), 90. 10.1007/s00018-024-05137-y

Lucien, F., Gustafson, D., Lenassi, M., Li, B., Teske, J. J., Boilard, E., von Hohenberg, K. C., Falcon-Perez, J. M., Gualerzi, A., Reale, A., Jones, J. C., Lasser, C., Lawson, C., Nazarenko, I., O’Driscoll, L., Pink, R., Siljander, P. R., Soekmadji, C., Hendrix, A.,…Nieuwland, R. (2023). MIBlood-EV: Minimal information to enhance the quality and reproducibility of blood extracellular vesicle research. J Extracell Vesicles, 12(12), e12385. 10.1002/jev2.12385

Ma, S. R., Xia, H. F., Gong, P., & Yu, Z. L. (2023). Red Blood Cell-Derived Extracellular Vesicles: An Overview of Current Research Progress, Challenges, and Opportunities. Biomedicines, 11(10). 10.3390/biomedicines11102798

Ma, W., Gil, H. J., Escobedo, N., Benito-Martin, A., Ximenez-Embun, P., Munoz, J., Peinado, H., Rockson, S. G., & Oliver, G. (2020). Platelet factor 4 is a biomarker for lymphatic-promoted disorders. JCI Insight, 5(13). 10.1172/jci.insight.135109

Malaguarnera, M., & Cabrera-Pastor, A. (2024). Emerging Role of Extracellular Vesicles as Biomarkers in Neurodegenerative Diseases and Their Clinical and Therapeutic Potential in Central Nervous System Pathologies. International Journal of Molecular Sciences, 25(18). 10.3390/ijms251810068

Manno, M., Bongiovanni, A., Margolis, L., Bergese, P., & Arosio, P. (2024). The physico-chemical landscape of extracellular vesicles. Nature Reviews Bioengineering, 3(1), 68–82. 10.1038/s44222-024-00255-5

Pan, S., Manabe, N., & Yamaguchi, Y. (2021). 3D Structures of IgA, IgM, and Components. Int J Mol Sci, 22(23). 10.3390/ijms222312776

Peter Klinken, S. (2002). Red blood cells. Int J Biochem Cell Biol, 34(12), 1513–1518. 10.1016/s1357-2725(02)00087-0

Rai, A., Huynh, K., Cross, J., Poh, Q. H., Fang, H., Claridge, B., Duong, T., Duarte, C., Shaw, J. E., Marwick, T. H., Meikle, P., & Greening, D. W. (2025). Multi-omics identify hallmark protein and lipid features of small extracellular vesicles circulating in human plasma. Nat Cell Biol, 27(12), 2167–2185. 10.1038/s41556-025-01795-7

Rappsilber, J., Ishihama, Y., & Mann, M. (2003). Stop and Go Extraction Tips for Matrix-Assisted Laser Desorption/Ionization, Nanoelectrospray, and LC/MS Sample Pretreatment in Proteomics. Analytical Chemistry, 75(3), 663–670. 10.1021/ac026117i

Ravenhill, B. J., Kanjee, U., Ahouidi, A., Nobre, L., Williamson, J., Goldberg, J. M., Antrobus, R., Dieye, T., Duraisingh, M. T., & Weekes, M. P. (2019). Quantitative comparative analysis of human erythrocyte surface proteins between individuals from two genetically distinct populations. Commun Biol, 2, 350. 10.1038/s42003-019-0596-y

Rayamajhi, S., Sipes, J., Tetlow, A. L., Saha, S., Bansal, A., & Godwin, A. K. (2024). Extracellular Vesicles as Liquid Biopsy Biomarkers across the Cancer Journey: From Early Detection to Recurrence. Clin Chem, 70(1), 206–219. 10.1093/clinchem/hvad176

Rios de Los Rios Resendiz, J., Herrmann-Sim, F., Wilkesmann, L., Helm, D., Schneider, M., Campione, G., Plugge, K., Greiner, G., Lazaro Garcia, L., Berker, J., Richter, K., Zielske, L., Hofmann, W. K., & Clemm von Hohenberg, K. (2025). A translational protocol optimizes the isolation of plasma-derived extracellular vesicle proteomics. Sci Rep, 15(1), 24292. 10.1038/s41598-025-08366-8

Shah, J. S., Soon, P. S., & Marsh, D. J. (2016). Comparison of Methodologies to Detect Low Levels of Hemolysis in Serum for Accurate Assessment of Serum microRNAs. PLoS One, 11(4), e0153200. 10.1371/journal.pone.0153200

Sidhom, K., Obi, P. O., & Saleem, A. (2020). A Review of Exosomal Isolation Methods: Is Size Exclusion Chromatography the Best Option? Int J Mol Sci, 21(18). 10.3390/ijms21186466

Singh, M., Tiwari, P. K., Kashyap, V., & Kumar, S. (2025). Proteomics of Extracellular Vesicles: Recent Updates, Challenges and Limitations. Proteomes, 13(1). 10.3390/proteomes13010012

Sódar, B. W., Kittel, Á., Pálóczi, K., Vukman, K. V., Osteikoetxea, X., Szabó-Taylor, K., Németh, A., Sperlágh, B., Baranyai, T., & Giricz, Z. (2016). Low-density lipoprotein mimics blood plasma-derived exosomes and microvesicles during isolation and detection. Scientific reports, 6(1), 1–12.

Sodar, B. W., Kittel, A., Paloczi, K., Vukman, K. V., Osteikoetxea, X., Szabo-Taylor, K., Nemeth, A., Sperlagh, B., Baranyai, T., Giricz, Z., Wiener, Z., Turiak, L., Drahos, L., Pallinger, E., Vekey, K., Ferdinandy, P., Falus, A., & Buzas, E. I. (2016). Low-density lipoprotein mimics blood plasma-derived exosomes and microvesicles during isolation and detection. Sci Rep, 6, 24316. 10.1038/srep24316

Soleymani, T., Chen, T. Y., Gonzalez-Kozlova, E., & Dogra, N. (2023). The human neurosecretome: extracellular vesicles and particles (EVPs) of the brain for intercellular communication, therapy, and liquid-biopsy applications. Front Mol Biosci, 10, 1156821. 10.3389/fmolb.2023.1156821

Stam, J., Bartel, S., Bischoff, R., & Wolters, J. C. (2021). Isolation of extracellular vesicles with combined enrichment methods. J Chromatogr B Analyt Technol Biomed Life Sci, 1169, 122604. 10.1016/j.jchromb.2021.122604

Stranska, R., Gysbrechts, L., Wouters, J., Vermeersch, P., Bloch, K., Dierickx, D., Andrei, G., & Snoeck, R. (2018). Comparison of membrane affinity-based method with size-exclusion chromatography for isolation of exosome-like vesicles from human plasma. J Transl Med, 16(1), 1. 10.1186/s12967-017-1374-6

Su, X., Junior, G. P. O., Marie, A. L., Gregus, M., Figueroa-Navedo, A., Ghiran, I. C., & Ivanov, A. R. (2024). Enhanced proteomic profiling of human plasma-derived extracellular vesicles through charge-based fractionation to advance biomarker discovery potential. J Extracell Vesicles, 13(12), e70024. 10.1002/jev2.70024

Suades, R., Padro, T., Vilahur, G., & Badimon, L. (2022). Platelet-released extracellular vesicles: the effects of thrombin activation. Cell Mol Life Sci, 79(3), 190. 10.1007/s00018-022-04222-4

Szklarczyk, D., Kirsch, R., Koutrouli, M., Nastou, K., Mehryary, F., Hachilif, R., Gable, A. L., Fang, T., Doncheva, N. T., Pyysalo, S., Bork, P., Jensen, L. J., & von Mering, C. (2023). The STRING database in 2023: protein-protein association networks and functional enrichment analyses for any sequenced genome of interest. Nucleic Acids Res, 51(D1), D638–D646. 10.1093/nar/gkac1000

Tang, D., Chen, M., Huang, X., Zhang, G., Zeng, L., Zhang, G., Wu, S., & Wang, Y. (2023). SRplot: A free online platform for data visualization and graphing. PLoS One, 18(11), e0294236. 10.1371/journal.pone.0294236

Ter-Ovanesyan, D., Gilboa, T., Budnik, B., Nikitina, A., Whiteman, S., Lazarovits, R., Trieu, W., Kalish, D., Church, G. M., & Walt, D. R. (2023a). Improved isolation of extracellular vesicles by removal of both free proteins and lipoproteins. eLife, 12, e86394. 10.7554/eLife.86394

Ter-Ovanesyan, D., Gilboa, T., Budnik, B., Nikitina, A., Whiteman, S., Lazarovits, R., Trieu, W., Kalish, D., Church, G. M., & Walt, D. R. (2023b). Improved isolation of extracellular vesicles by removal of both free proteins and lipoproteins. Elife, 12. 10.7554/eLife.86394

Thery, C., Amigorena, S., Raposo, G., & Clayton, A. (2006). Isolation and characterization of exosomes from cell culture supernatants and biological fluids. *Curr Protoc Cell Biol*, Chapter 3, Unit 3 22. 10.1002/0471143030.cb0322s30

Thery, C., Witwer, K. W., Aikawa, E., Alcaraz, M. J., Anderson, J. D., Andriantsitohaina, R., Antoniou, A., Arab, T., Archer, F., Atkin-Smith, G. K., Ayre, D. C., Bach, J. M., Bachurski, D., Baharvand, H., Balaj, L., Baldacchino, S., Bauer, N. N., Baxter, A. A., Bebawy, M.,…Zuba-Surma, E. K. (2018). Minimal information for studies of extracellular vesicles 2018 (MISEV2018): a position statement of the International Society for Extracellular Vesicles and update of the MISEV2014 guidelines. J Extracell Vesicles, 7(1), 1535750. 10.1080/20013078.2018.1535750

Vallejo, M. C., Sarkar, S., Elliott, E. C., Henry, H. R., Powell, S. M., Diaz Ludovico, I., You, Y., Huang, F., Payne, S. H., Ramanadham, S., Sims, E. K., Metz, T. O., Mirmira, R. G., & Nakayasu, E. S. (2023). A proteomic meta-analysis refinement of plasma extracellular vesicles. Sci Data, 10(1), 837. 10.1038/s41597-023-02748-1

Van Deun, J., Jo, A., Li, H., Lin, H. Y., Weissleder, R., Im, H., & Lee, H. (2020). Integrated Dual-Mode Chromatography to Enrich Extracellular Vesicles from Plasma. Adv Biosyst, 4(12), e1900310. 10.1002/adbi.201900310

van Niel, G., D’Angelo, G., & Raposo, G. (2018). Shedding light on the cell biology of extracellular vesicles. Nat Rev Mol Cell Biol, 19(4), 213–228. 10.1038/nrm.2017.125

Vanderboom, P. M., Dasari, S., Ruegsegger, G. N., Pataky, M. W., Lucien, F., Heppelmann, C. J., Lanza, I. R., & Nair, K. S. (2021). A size-exclusion-based approach for purifying extracellular vesicles from human plasma. Cell Rep Methods, 1(3). 10.1016/j.crmeth.2021.100055

Veerman, R. E., Teeuwen, L., Czarnewski, P., Gucluler Akpinar, G., Sandberg, A., Cao, X., Pernemalm, M., Orre, L. M., Gabrielsson, S., & Eldh, M. (2021). Molecular evaluation of five different isolation methods for extracellular vesicles reveals different clinical applicability and subcellular origin. J Extracell Vesicles, 10(9), e12128. 10.1002/jev2.12128

Venturella, M., Carpi, F. M., & Zocco, D. (2019). Standardization of blood collection and processing for the diagnostic use of extracellular vesicles. Current Pathobiology Reports, 7(1), 1–8.

Wan Azman, W. N., Omar, J., Koon, T. S., & Tuan Ismail, T. S. (2019). Hemolyzed Specimens: Major Challenge for Identifying and Rejecting Specimens in Clinical Laboratories. Oman Med J, 34(2), 94–98. 10.5001/omj.2019.19

Welsh, J. A., Goberdhan, D. C. I., O’Driscoll, L., Buzas, E. I., Blenkiron, C., Bussolati, B., Cai, H., Di Vizio, D., Driedonks, T. A. P., Erdbrugger, U., Falcon-Perez, J. M., Fu, Q. L., Hill, A. F., Lenassi, M., Lim, S. K., Mahoney, M. G., Mohanty, S., Moller, A., Nieuwland, R.,…Witwer, K. W. (2024). Minimal information for studies of extracellular vesicles (MISEV2023): From basic to advanced approaches. J Extracell Vesicles, 13(2), e12404. 10.1002/jev2.12404

Wiersema, A. F., Rennenberg, A., Smith, G., Varderidou-Minasian, S., & Pasterkamp, R. J. (2024). Shared and distinct changes in the molecular cargo of extracellular vesicles in different neurodegenerative diseases. Cell Mol Life Sci, 81(1), 479. 10.1007/s00018-024-05522-7

Wu, C. C., Tsantilas, K. A., Park, J., Plubell, D., Sanders, J. A., Naicker, P., Govender, I., Buthelezi, S., Stoychev, S., Jordaan, J., Merrihew, G., Huang, E., Parker, E. D., Riffle, M., Hoofnagle, A. N., Noble, W. S., Poston, K. L., Montine, T. J., & MacCoss, M. J. (2025). Enrichment of extracellular vesicles using Mag-Net for the analysis of the plasma proteome. Nat Commun, 16(1), 5447. 10.1038/s41467-025-60595-7

Yin, H., Xie, J., Xing, S., Lu, X., Yu, Y., Ren, Y., Tao, J., He, G., Zhang, L., Yuan, X., Yang, Z., & Huang, Z. (2024a). Machine learning-based analysis identifies and validates serum exosomal proteomic signatures for the diagnosis of colorectal cancer. Cell Rep Med, 5(8), 101689. 10.1016/j.xcrm.2024.101689

Yin, V., Desligniere, E., Mokiem, N., Gazi, I., Lood, R., de Haas, C. J. C., Rooijakkers, S. H. M., & Heck, A. J. R. (2024b). Not All Arms of IgM Are Equal: Following Hinge-Directed Cleavage by Online Native SEC-Orbitrap-Based CDMS. J Am Soc Mass Spectrom, 35(6), 1320–1329. 10.1021/jasms.4c00094

