## Supplementary material for "Combining Anion Exchange and Size Exclusion Chromatography for Extracellular Vesicle Enrichment from Small Volumes of Human and Mouse Plasma for Quantitative Proteomics": Graphical Abstract

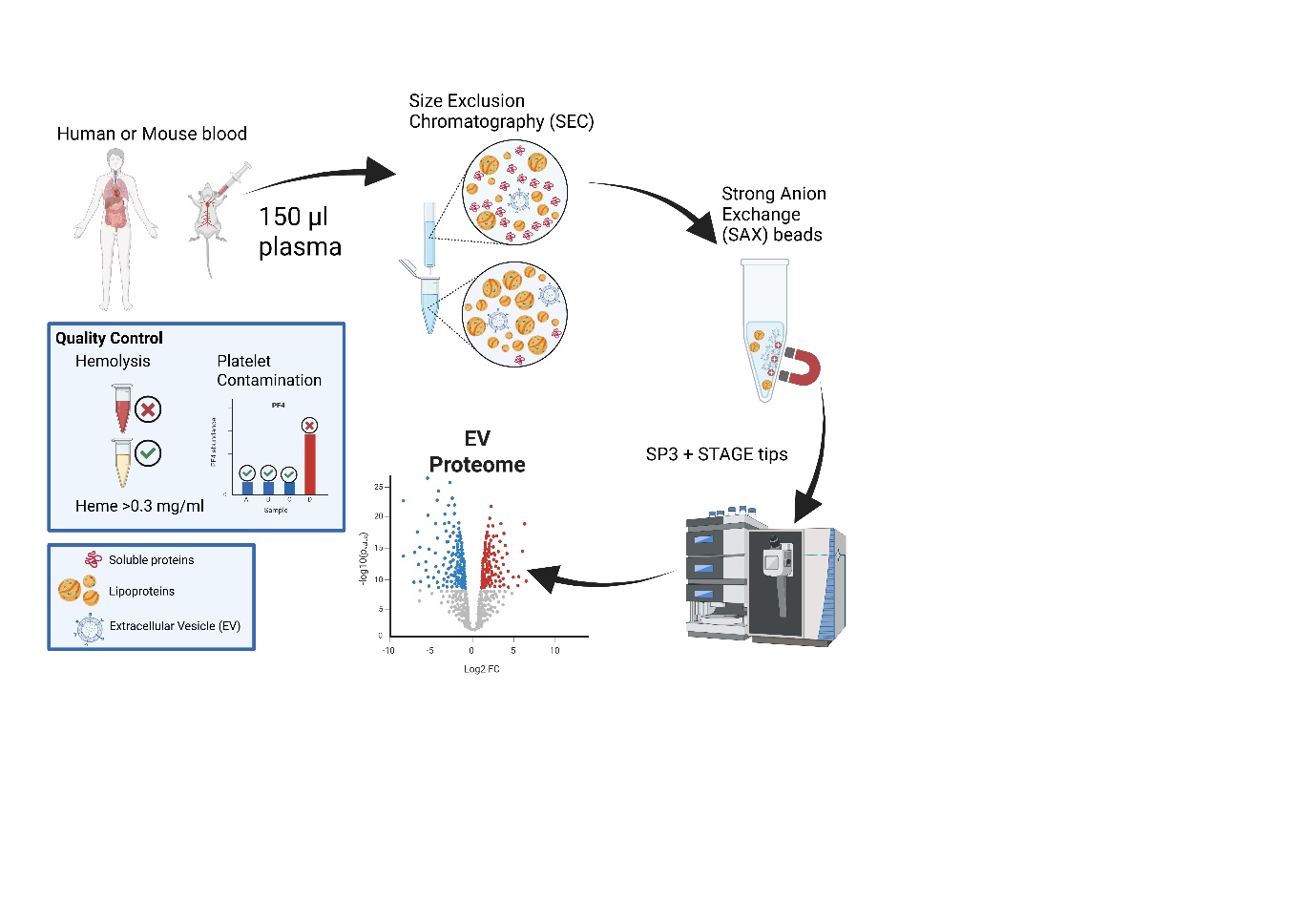

Using 150 µl plasma, the SEC-SAX workflow improves EV proteome depth by reducing ApoB lipoproteins and enriching EVs for Orbitrap analysis following SP3 and STAGE-tip processing. QC measures include excluding samples with hemolysis ≥0.3 mg/mL heme and identifying platelet contamination via PF4. The method was systematically optimized across bead amount, digestion, and cleanup steps, and validated in both human and mouse plasma.
